# Dendritic cell targeting in lymph nodes with engineered modular adapters improves HAdV5 and HC-HAdV5 tumor vaccination by co-secretion of IL-2v and IL-21

**DOI:** 10.1101/2024.04.27.591433

**Authors:** Fabian Weiss, Jonas Kolibius, Patrick C. Freitag, Felix Gantenbein, Anja Kipar, Andreas Plückthun

**Author notes:** Denotes co-second author (Jonas Kolibius, Patrick C. Freitag). Correspondence should be addressed to A.P.

## Abstract

Adenoviral vectors demonstrate encouraging clinical outcomes for B- and T-cell vaccines. With such approaches, multiple payloads can be delivered, beyond the antigen itself. Nevertheless, the human adenoviral vector serotype C5 (HAdV5) exhibits limited transduction efficiency to dendritic cells (DC), therefore necessitating very high viral loads. Targeting antigen-presenting cells (APC) has remained challenging. To solve this problem, we developed a versatile platform that employs modular retargeting adapters to enhance transduction of specific cell types, including challenging host cells. By rational design, we constructed a dual-adapter for DC-SIGN and CD11c and demonstrate successful targeting of HAdV5 to human and murine DCs. Our *in vivo* characterization highlights improved and specific transduction of DCs in draining lymph nodes. Moreover, a tumor vaccination study showcases the advantageous co-expression of T cell stimulatory cytokines (IL-2v or IL-21) locally in lymph nodes alongside a potent tumor antigen. Lymph node-directed gene therapy at significantly reduced vector loads circumvents potential systemic toxicity of stimulating payloads. Our proposed low-dosage DC-targeted vaccine offers an effective solution for patients and also minimizes potential adenovirus-related side-effects. The robust immunogenicity of HC-HAdV5, with its large coding capacity (37 kbp DNA), opens up exciting possibilities for future therapeutic combination strategies.

## Introduction

Cancer remains one of the major unsolved global health concerns alongside chronic heart disease. Decades of research have increased our understanding of this disease from the initial discovery of p53 to the definition of the hallmarks of cancer,^1^ revealing the multifaceted complexity of this disease. Over the past few decades, immunotherapy has emerged as a promising approach to fight cancer by stimulating the immune system to identify and eliminate neoplastic cells. Immunotherapy represents by now a main treatment modality besides chemo- and radiotherapy and surgical removal of cancerous tissue. Consequently, a multitude of checkpoint-blockade therapies and T cell engagers have entered clinical trials, and many of them were approved for clinical application.^2,3^

Tumor vaccination is yet another immunotherapy approach to prime the patient’s immune system to eliminate cancer cells by supporting and complementing the cancer immunity (CI) cycle.^4^ Initially considered clinically insignificant, major progress has been made in the last ten years through recent advances in neoantigen research in conjunction with enhanced understanding of antigen processing and powerful gene delivery methods.^5,6^ Tumor vaccination enhances the generation of active cytotoxic T cells with matching T cell receptors (TCRs) against defined cancer antigens. This approach entails administration of tumor-associated antigens (TAAs) or neoantigens obtained from the patient’s own tumor and uptake by professional antigen presenting cells (APCs) to initiate an antigen-specific immune response.

Dendritic cells (DCs) act as key conductors of the CI cycle by serving as APC, which can elicit and regulate immune responses. Their unique ability to capture, process and present antigens to T cells renders them crucial in shaping adaptive immunity. Traditional DC vaccination strategies involve the isolation and *ex vivo* manipulation of DCs to load them with tumor-derived peptide antigens prior to their reintroduction into the patient’s body.^7^ However, *ex vivo* DC vaccination faces several limitations that have dampened its clinical success. These challenges consist of a short *ex vivo* half-life, loss of antigen, necessary clinical standardization (requiring suitable cell therapy facilities), and thus high health care costs. As an approach to circumvent these obstacles, we propose the utilization of DC-targeted *in vivo* gene-based therapies for the delivery of multiple payloads.

Non-integrating human adenoviruses are among the most commonly used vectors in gene therapy.^8,9^ They demonstrate substantial promise for vaccine vector development due to their capacity to evoke T and B cell responses while exhibiting relatively low pathogenicity in humans. The best clinical examples so far are the adenoviral COVID-19 vaccines Vaxzevria (AstraZeneca), Sputnik V (Gamaleya Research Institute, Acellena Contract Drug Research and Development), JNJ-78436735 (Janssen) or Convidecia (CanSino Biologics), which are based on human and non-human AdV serotypes.^10^ The potential of adenoviruses to induce maturation of DCs, thereby acting as vaccination boosters, makes them popular for vaccination approaches.^10^ Pathogen-associated molecular patterns (PAMPs) trigger the innate response, causing in the DCs the upregulation and activation of co-receptors CD40, CD80, and CD86, which are important for T cell activation.^11-13^ The subsequent immunological synapse at the interface between APC and T cells results in priming and proliferation of specific T cells with matching TCR to the presented antigen (pMHC).

Over 100 human adenovirus types have been identified so far, the most extensively studied and widely utilized being the human adenovirus serotype C5 (HAdV5).^8^ For clinical applications, 1^st^ generation vectors with a maximum coding capacity of 7.5 kb DNA have been constructed by deleting gene regions E1 and E3 from this non-enveloped double-stranded DNA virus to render it replication-deficient.^8,14^ In contrast, the gutless adenovirus or high-capacity adenovirus (HC-HAdV5) represents a next generation vector that is developed by deleting all viral coding sequences except the inverted terminal repeats (ITRs) and the packaging signal. Due to its large transgene capacity of up to 37 kb, HC-HAdV5 is particularly appropriate for delivering large and multiple transgenes.^15,16^ While HC-HAdV5 is currently not extensively utilized in vaccination studies, primarily because of its more intricate production process, significant advances in fast and high-purity large scale production have lately been made.^17-19^ Despite the deletion of all adenoviral genes and hence the lack of adenoviral protein expression, HC-HAdV5-vectors still elicit robust innate and adaptive immune responses.^20^ The host cell detects transduction by HC-HAdV5-vectors mainly through recognition of unmethylated CpG sequences by TLR9^21^ or the presence of cytosolic dsDNA (via the c-GAS-STING pathway^22^). Moreover, TLR2 and/or TLR4 recognize the viral capsid on the surface of APCs through viral particle complexes with coagulation factor X (FX) or lactoferrin.^23-26^

The natural cellular entry of HAdV5 vectors occurs through contact of the homotrimeric fiber-knob protein with the coxsackie and adenovirus receptor (CAR),^27^ followed by interaction between the RGD-motif in the penton base and a_v_β_3_-or a_v_β_5_-integrins on the cell membrane.^28^ Subsequently, the virus is internalized by clathrin-mediated endocytosis,^29^ followed by fiber shedding, endosomal escape, and microtubule-based trafficking to the nucleus. Upon docking to the nuclear pore complex, the adenoviral DNA is translocated from the capsid into the nucleus.^30^

Intravenously administered HAdV5 has a strong liver tropism, attributed to the interplay between coagulation factor X (FX), complement factors, and non-neutralizing antibodies with epitopes on the capsid’s hypervariable (HVR) loops. This results in its effective clearance by hepatocytes and Kupffer cells.^31,32^ Vaccination vectors are usually administered intramuscularly to patients, but the precise mode of delivery of the encoded antigen to APCs and the subsequent presentation in the lymph nodes to T cells is uncertain. Since human adenoviral vectors can only marginally transduce muscle cells,^33,34^ the direct gene-transfer to DCs located in the lymph nodes may constitute a more efficient vaccination alternative, but it remains a challenge. DCs do not express CAR and only low levels of RGD-binding integrins.^35,36^ Additionally, they are a rare cell type, which makes them difficult to target as they are only enriched to a maximum of 1% in spleen, lymph nodes or Peripheral Blood Mononuclear Cells (PBMCs). Consequently, the question remains open how to efficiently redirect adenoviral vectors to DCs and facilitate potent gene delivery of the encoded antigen, along with genes for auxiliary factors, while avoiding off-targeting to other susceptible cells that are much more abundant compared to the target cell population.

Our lab developed a gene-delivery system based on HAdV5 that targets the vector to specific cell types while simultaneously inhibiting the natural tropism.^37^ Our technology has allowed for direct transduction of T cells,^38^ fibroblasts,^39^ or cancer cells^19,40^ *in vivo,* while drastically reducing the considerable viral tropism to the liver by using shielded vectors.^41^ We have achieved this by using stable trimerized protein clamps that bind with high avidity to the HAdV5 knob, covering the CAR epitope and effectively reducing natural transduction via CAR. Transduction can now be redirected to surface cell markers through fused retargeting moieties such as DARPins, scFvs, peptides or small molecules^42^.

In this study, we are expanding our portfolio by introducing dual-adapters to both DC-SIGN (CD209) and CD11c, which led to increased transduction of human and murine DCs while maintaining their specificity *in vivo*. These results propose a promising strategy to strengthen the effectiveness of tumor vaccination via directed gene delivery targeting lymph nodes and by encoding auxiliary factors. We report here the autocrine and paracrine immune stimulation of IL-2v or IL-21, co-expressed by the same vaccine vector, which allows a potent vaccination effect at an exceptionally low dosage of administered HC-HAdV5. This approach has significant potential to advance the field of combination immunotherapy by enhancing the precision of effective DC-based gene therapies, with the additional potential of significantly reducing costs of cell-based therapies.

## Results

### DC-SIGN was identified as best target for mono-adapters mediating HAdV5 transduction

Adapter-mediated targeting of a specific cell type requires a surface marker that can be used to redirect the binding of an adenoviral vector^37-42^. It is necessary for this surface marker to be expressed at sufficient levels to make the cell susceptible for transduction. Additionally, the surface protein also needs to be expressed in a cell-specific manner to prevent off-targeting *in vivo*. Therefore, we screened multiple retargeting moieties (scFv, peptides or viral glycoproteins) that bind to DC-SIGN (DC209)^43^, CD11c^44,45^, DEC205 (CD205)^45,46^, CD206^47^ and CD40^48^, using the sequences from the indicated publications. We selected these targets, since they are described as surface markers for immature DCs, or these targets have already been used to direct antigens or viruses to DCs. For our preliminary study we used the cell line DC2.4^49^, an immortalized and stable DC line transduced with GM-CSF, isolated from the bone marrow of C57BL/6 mice. Immature DC2.4 cells express low levels of CD40, CD206 and DEC-205, whereas DC-SIGN and CD11c are moderately expressed (Figure S1), reflecting the expression pattern of primary immature DCs of mouse and human origin.^50,51^ To target HAdV5 to DCs, we first employed an adapter system comprising an N-terminal retargeting moiety with a single specificity, fused via a flexible linker to a DARPin (1D3) which binds to the fiber knob and covers the CAR epitope, i.e. of a similar format as described perviously^37^. We term this a mono-adapter. The C-terminal SHP trimerization domain (derived from lambdoid phage 21) forms a very stable trimerized protein clamp around the HAdV5 fiber knob (Figure 1A).^37^

**Figure 1:**
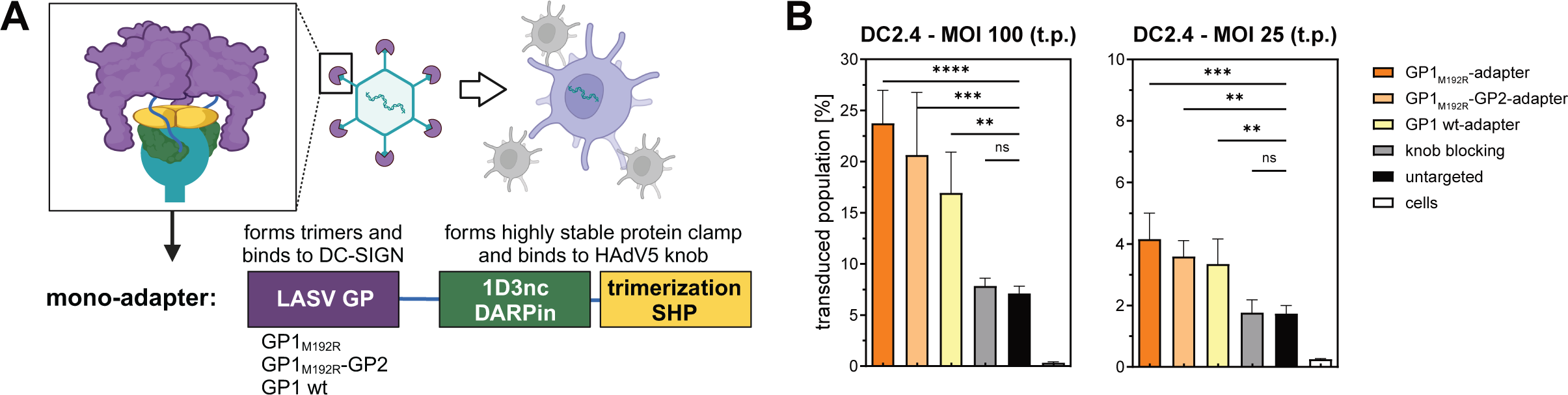
Retargeting module GP_LASV_ mediates HAdV5^ΔHVR7^ transduction of murine dendritic cells via DC-SIGN control. (A) Schematic representation of a trimeric retargeting adapter. The retargeting module (violet), is fused by a flexible linker to the knob binding DARPin 1D3nc (green). The C-terminal trimerizing protein SHP (yellow) leads to the formation of a stable clamp formed around the fiber knob (cyan) and blocks the epitope binding to CAR (coxsackievirus and adenovirus receptor). The retargeting domain, in this case GP_LASV_, redirects binding to a selected surface marker (e.g. DC-SIGN) and can mediate transduction of dendritic cells. The transduced cell (violet) produces a reporter protein. GP_LASV_ is trimeric. (B) Flow cytometry analysis of a transduction experiment with HAdV5^ΔHVR7^, expressing tdTomato as reporter protein. The tdTomato^+^ population of DC2.4 cells is depicted. The glycoprotein of the Lassa virus (LASV) mediates higher transduction via the surface marker DC-SIGN in the mono-adapter format in comparison to the untargeted control group. Three variations of GP_LASV_ were tested as retargeting domain: GP comprising solely the GP1 wild-type domain (**GP1 wt-adapter**) (amino acids 59-237); a full-length GP carrying an M192R mutation (**GP1_M192R_-GP2-adapter**) (amino acids 59-424); and a fragment of GP consisting only of the GP1 domain also carrying the M192R mutation (**GP1_M192R_-adapter**) (amino acids 59-237). The mutation M192R depletes binding of GP1 to LAMP1. A non-binding adapter (E2_5) was used as negative control for knob blocking in order to block natural transduction via CAR. t.p: transducing particles. Statistics: Representative data of two independent experiments shown. P-value determined using one-way ANOVA with Dunnett’s test for multiple comparisons to untargeted control. *p < 0.05, **p < 0.01, ***p < 0.001, ****p < 0.0001. Bar graphs represent mean ± SD, n = 4.

To evaluate transduction via the DC targets mentioned above, a diverse set of retargeting mono-adapters consisting of scFvs, peptides or viral glycoproteins was recombinantly produced in mammalian cell culture and purified by immobilized metal ion affinity chromatography (IMAC). SDS-PAGE confirmed the purity and the expected molecular weights (Figure S2). To assess the capability of the individual adapters to mediate transduction of DCs, we first complexed a tdTomato-encoding HAdV5 with the respective adapter and then added the modified vector to the DC2.4 target cells. Adapter-mediated transduction efficiency was monitored by quantifying the percentage of transduced cells. Flow cytometry data (Figure S2D) from 25 adapters indicated successful transduction events for six binders to the leukocyte integrin CD11c (integrin α-X)^52^ and one binder to the endocytic receptor DEC-205^53^. In contrast, CD40^54^- and CD206^55^-binding adapters mediated no significant transduction. It should be highlighted that those surface markers are only expressed at low levels on dendritic cells and therefore generally do not represent an ideal target. The three most positive hits were achieved using the Lassa virus (LASV) glycoprotein and its derivatives targeting DC-SIGN^56-58^. Here we describe the rationale of the DC-SIGN-binding modules in more detail.

DC-SIGN (or CD209) is a transmembrane C-type lectin receptor. The tetrameric receptor with its characteristic tandem-repeats binds with the extracellular C-terminal domain to high-mannose carbohydrates.^56-58^ This is one of the mechanisms by which DCs recognize microbial pathogens or viruses on the cell surface. Lassa virus is capable to naturally transduce human DCs and monocytes. It was shown that the glycoprotein domain 1 (GP1) of LASV binds to DC-SIGN (or α-dystroglycan) and thereby mediates natural infection of DCs.^43^ The GP of LASV, with a total of eleven N-linked glycans, consists of two main domains, GP1 (amino acids 59-259) and GP2 (amino acids 260-432). In an acidified endosome, the internalized LASV particle dissociates from DC-SIGN and then attaches to LAMP1. The interaction between DC-SIGN and LAMP1 was previously studied.^59,60^

Therefore, we explored how LAMP1 interactions with GP_LASV_ of the retargeting adapter influences transduction mediated by a retargeted HAdV5 particle, as this interaction might redirect the import pathway. To examine this interaction, we introduced the point mutation M192R in GP1, which results in blocked binding to LAMP1.^60^ In total, three versions of GP were tested (GP1_wt_, GP1_M192R_ and GP1_M192R_-GP2) to assess their capacity for transducing DCs (Figure 1A). For this purpose, we designed different adapters carrying as retargeting module either one of the three GP versions. All three samples mediated higher transduction in comparison to the untargeted control for two tested MOIs (multiplicity of infection). They also clearly exceeded the rates of other mono-adapters (Figure S2). Importantly, the two adapters with the retargeting domain containing the M192R mutation (GP1_M192R_ and GP1_M192R_-GP2) that are not binding to LAMP1 showed higher transduction efficiency than GP1_wt_. (Figure 1B). This effect was especially prominent at a high vector load (MOI 100), while the differences were less evident at a low vector load (MOI 25). In conclusion, the GP1_M192R_-adapter targeting DC-SIGN was the most effective mono-adapter tested in the first screening round, further suggesting that a transfer from DC-SIGN to LAMP1 binding was not favorable for HAdV5 transduction.

### Optimized dual-adapter engaging DC-SIGN and CD11c increases transduction of murine and human primary DCs

To further improve the adapter performance, we developed the concept of dual-adapters. A dual-adapter contains two fused retargeting moieties in tandem, connected by a flexible linker. This design enables the adenoviral particle to bind to either of the targeted surface markers, or potentially to both simultaneously, thereby increasing avidity (Figure 2A and S3A). Initially, we investigated the feasibility of rescuing those retargeting moieties that lacked sufficient functionality in mono-adapter form. Hence, we combined DEC205 and CD11c targeting units in different N- to C-terminal arrangements connected with variable linker lengths. We determined the HAdV5 transduction efficiencies with an encoded fluorescent reporter protein (tdTomato). In comparison to mono-module DARPin adapters, increased transduction efficiencies were observed for the majority of dual-module adapters tested (Figure S3). Encouraged by this trend, we then generated multiple combinations of dual-adapters using the LASV GP1_M192R_ domain, which had provided excellent results already as a mono-targeting unit. Since GP1 is a trimer itself, it was situated in proximity to the trimeric knob-clamp (DARPin 1D3nc-SHP), so that it can form the native assembly. The flexible retargeting unit was then fused to the N-terminus.

**Figure 2:**
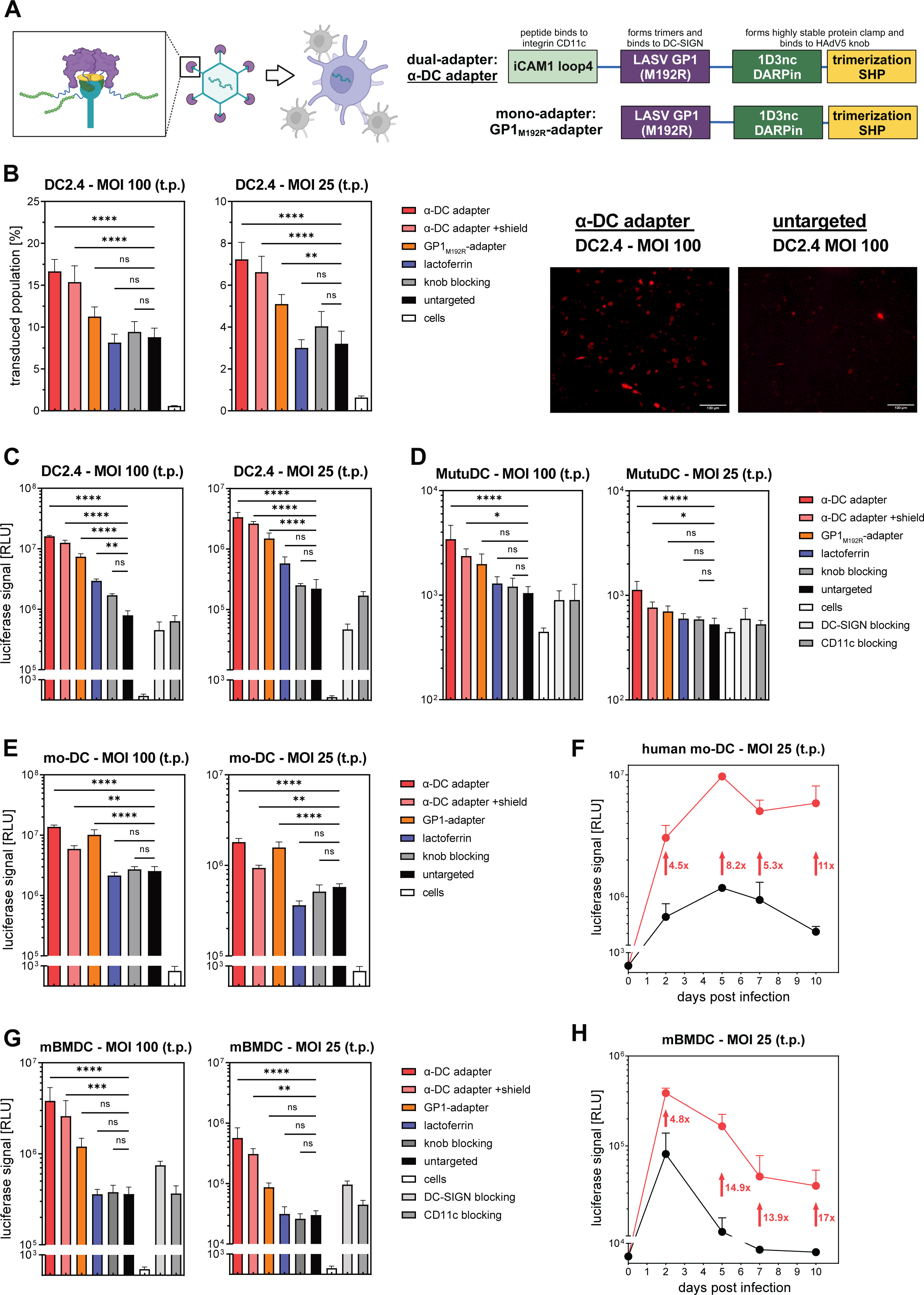
Dual-adapter (α-DC adapter) mediates HAdV5^ΔHVR7^ transduction of human and murine dendritic cells via DC-SIGN and CD11c. (A) Schematic representation of a dual-adapter. In contrast to the mono-adapter (GP1_M192R_-adapter), the dual-adapter (the preferred design is termed α-DC adapter) consists of two retargeting modules which are connected by a flexible linker. The N-terminal module targets the integrin CD11c with a peptide derived from the iCAM1 loop4, followed by the DC-SIGN-targeting module, LASV GP1_M192R_. The dual-adapter forms a stable trimeric clamp around the fiber knob. (B-H) Determination of transduction efficiency by intracellular expression of transgenes measured by flow cytometry and fluorescent microcopy or luminescence. *α-DC adapter,* retargeting HAdV5^ΔHVR7^ with dual-adapter leads to a strong increase of transduction efficiency at various MOI (100 or 25) compared to untargeted control. *α-DC adapter +shield*, retargeted HAdV5^ΔHVR7^ with dual-adapter and trimeric hexon-shielding protein. *GP1_M192R_-adapter*, retargeting HAdV5^ΔHVR7^ with mono-adapter as control. *Lactoferrin*, coating HAdV5^ΔHVR7^ with hexon-binding lactoferrin (1 µM) as control. *Knob-blocking* and *Untargeted*, as negative controls were used HAdV5^ΔHVR7^ with blocked knob (E2_5 blocking adapter), and an untargeted control (only HAdV5^ΔHVR7^) was tested to determine the natural transduction rate. *DC-SIGN blocking*, transduction of targeted (+α-DC adapter) HAdV5^ΔHVR7^ was blocked by antibodies binding to DC-SIGN. *CD11c blocking*, transduction of targeted (+α-DC adapter) HAdV5^ΔHVR7^ was blocked by antibodies binding to CD11c. (B) Murine cell line DC2.4 transduced by HAdV5^ΔHVR7^ encoding tdTomato or (C) ffLuciferase. (D) Cell line MutuDC transduced by HAdV5^ΔHVR7^ encoding ffLuciferase. (E) Primary human monocyte-derived dendritic cells (human mo-DC) transduced by HAdV5^ΔHVR7^ encoding ffLuciferase. (F) Transduced primary human monocyte-derived dendritic cells (human mo-DC) express the transgene ffLuciferase over a long time course for at least 10 days. Red = Retargeted HAdV5^ΔHVR7^ with α-DC adapter; Black = Untargeted HAdV5^ΔHVR7^ (G) Murine bone marrow-derived dendritic cells (mBMDC) transduced by HAdV5^ΔHVR7^ encoding ffLuciferase. (H) Transduced primary murine bone marrow-derived dendritic cells (mBMDC) express the transgene ffLuciferase over a long time course for at least 10 days. Red = retargeted HAdV5^ΔHVR7^ with α-DC adapter; Black = untargeted HAdV5^ΔHVR7^. t.p: transducing particles. Statistics: Representative data of two or three independent experiments shown. P-value determined using one-way ANOVA with Dunnett’s test for multiple comparisons to untargeted control. *p < 0.05, **p < 0.01, ***p < 0.001, ****p < 0.0001. Bar graphs represent mean ± SD, n = 4. Microscopic scale bars 130 µm

We concentrate here on our lead dual-adapter (henceforth termed the α-DC adapter), featuring the highest transduction efficiencies: This α-DC adapter is comprised of the loop 4 of iCAM1, i.e. an anti-CD11c moiety,^61,62^ linked to the described LASV GP1_M192R_ (Figure 2A). This construct was expressed in mammalian cell culture, and after affinity chromatography, it was polished by size exclusion chromatography (SEC) for high monomeric content. Purity was confirmed using reducing SDS-PAGE, and binding to human and murine DC-SIGN was confirmed by ELISA (Figure S4).

The α-DC adapter demonstrated a robust increase in mediated HAdV5 transduction of DC2.4 cells (MOI 100: 17%; MOI 25: 7% transduced cells) compared to the untargeted vector control (MOI 100: 8%; MOI 25: 3% transduced cells) (Figure 2B). A similar observation was made using an orthogonal readout when testing a HAdV5 reporter vector encoding ffLuciferase: The amount of total reporter protein was increased 15 to 20-fold in DC2.4 cells (MOI 100 and 25) (Figure 2C). To verify these findings, we also measured transduction in an alternative mouse DC cell line expressing CD11c and DC-SIGN. For this purpose, we tested the splenic MutuDC cell line,^49,63^ which was inefficiently transduced by untargeted HAdV5. Indeed, upon complexation with α-DC adapter, the ffLuciferase activity was amplified by 5 to 8-fold (Figure 2D). Notably, the transduction level remained nearly constant when we added the anti-hexon binding protein shield^41^ (Figures 2B-D). This indicated that binding of the adapter to CD11c and DC-SIGN induced endocytosis, rather than that uptake occurs by an hexon-mediated entry mechanism. Additionally, we could demonstrate the clear benefit of dual-targeting with a side-by-side comparison of the α-DC adapter versus the GP1_M192R_-adapter: When the CD11c binding moiety was present (α-DC adapter) the tdTomato^+^ population increased in DC2.4 cells from 11% to 17% (at MOI 100), and from 5% to 8% (at MOI 25), respectively, compared to the GP1_M192R_-adapter (Figure 2B). A similar effect was also observed in experiments with HAdV5_ffLuc_, which resulted in a 2 to 3-fold increase in transduction signal in DC2.4 cells and MutuDC cells (Figures 2C and 2D).

Building on our previous results, we sought to validate our platform with human and murine primary dendritic cells. Briefly, we found an enhanced transduction by 5 to 11-fold in human monocyte-derived DCs (mo-DCs), utilizing the α-DC adapter compared to the untargeted control, depending on the MOI tested and the time point post transduction (Figures 2E and 2F). This trend was also observable in murine dendritic cells derived from the bone marrow (mBMDC), augmenting transduction efficiency by 5 to 19-fold (depending on the MOI tested and the time point post transduction) (Figures 2G and 2H). These findings also confirmed the cross-specificity of the α-DC adapter for human and mouse, which was expected based on the sequence similarity of the entry receptors (Figure S5). Our rationale of CD11c/DC-SIGN-induced internalization was further supported by the blocking of those surface markers with antibodies, whereby the transduction signal returned to the level of the untargeted control, confirming the adapter-mediated entry mechanism (Figure 2C, 2D and 2G). The adapter-mediated transduction was also confirmed for two alternative human donors at low MOI (Figure S6).

Upon deletion of the CD11c-retargeting moiety, the retargeting effects experienced a slight drop in human DCs by ⁓1.3-fold (α-DC adapter versus GP1_M192R_-adapter), whereas the effect was significantly stronger in murine DCs, resulting in 3 to 7-fold transduction loss (Figure 2E and 2G). It is important to note that the CD11c retargeting moiety, a 15-amino acid peptide, was derived from murine iCAM1 to achieve optimal effects in a mouse *in vivo* model. While the iCAM1 loop4 in humans shares high sequence similarity (Figure S5), we hypothesize that replacing the murine iCAM1 loop4 with its human counterpart may enhance the retargeting effect on human CD11c. However, the initial goal of this study was to test the approach in mice *in vivo* (see below).

Interestingly, we observed a significant decrease of α-DC adapter-mediated transduction signal by adding the hexon-binding shield protein^41^ (α-DC adapter vs α-DC adapter +shield) (Figures 2E and 2G). Although not observed in the particular murine cell lines that were tested, this effect was evident in primary DCs of both mice and human origin. We assume that this decrease resulted from sterically blocked interactions between surface markers of primary DCs and the HAdV5 hexon. While we did not further identify the specific marker responsible, we hypothesize that it might belong to the family of scavenger receptors. Particularly the scavenger receptor SR-A6 (MARCO) was previously described to facilitate adenoviral entry in macrophages^64^ and shares high sequence similarity with SR-A5 of DCs (Human Protein Atlas)^65^. Thus, SR-A5 might serve, via its interaction with HAdV5 hexon, as an alternative entry receptor in DCs. In contrast, the blocking of CAR-binding to the HAdV5 knob remained at the level of the untargeted control, highlighting that the untargeted entry in the CAR_low_ primary DCs is not CAR-dependent (Figures 2E and 2G). As an additional reference, we always included lactoferrin coating (1 µM) of the HAdV5 vector, since lactoferrin had previously been used as a DC transduction agent, which is not cell-specific, however. Our results showed that our developed α-DC adapter outcompeted the lactoferrin-mediated uptake by a factor ≥4 (Figures 2B-E and 2G).

The expression duration of transgenes remains a critical measure in gene therapy. We assessed ffLuciferase expression for 10 days post HAdV5 transduction (Figures 2F and 2H). Our results indicated that using the α-DC adapter for targeted transduction provided a long-term advantage regarding transgene expression levels over the untargeted control in primary DCs, while preserving the typical morphology of a healthy DC.

### α**-**DC adapter boosts targeting and transgene expression of DCs in lymph nodes without off-targeting

In order to achieve the best vaccination responses, it is vital to administer vaccines via the appropriate route. For the stimulation of immune responses, vaccines are typically injected subcutaneously or intramuscularly, and in some instances even intravenously. In our case, the objective was to administer the targeted vector in a way that maximizes transduction of DCs, which should ideally then migrate to the lymph nodes and interact there with lymphocytes to generate the desired immune responses. The intramuscular route of application should generally be avoided for mice, as mouse muscles are small and the injection can be painful for the animal^66^. In addition, it is inefficient to target DCs directly in muscular tissue since there are no DCs present in the healthy skeletal musculature and DC-like cells only migrate into this environment in response to injury and viral challenge.^67^ On the other hand, upon intravenous (*i.v.*) administration HAdV5 is rapidly cleared by the liver as a consequence of its prominent hepatotropism. Although shielding the HAdV5 particles partially overcame this tropism,^41^ the rescued particles failed to adequately drain into lymph nodes after *i.v.* tail vein administration (according to our pilot study, data not shown).

To further improve HAdV5 accumulation in the draining lymph nodes, we investigated subcutaneous administration of targeted viral vectors. We followed a previously published approach for rats and mice and placed the injection into the subcutis at the hock of the mice.^68,69^ A preliminary study confirmed that the vector particles successfully transduced cells in the draining lymph nodes, similar to transduction after subcutaneous injection into the foot pad (F. Weiss and F. Gantenbein, unpublished data). Hence we performed a study with HAdV5 retargeted with an α-DC adapter and an untargeted HAdV5 control group using injection in the hock (Figure 3A). We observed a 5-fold increase in the encoded reporter protein ffLuciferase activity in popliteal and inguinal lymph nodes when using the α-DC adapter over the untargeted control, and we did not detect any off-targeted ffLuciferase activity in liver and spleen (Figure 3C). The histological and immunohistochemical specimens were examined by a veterinary pathologist (A.K.), who was blinded to the treatment of the animals and the targeted cell type. Immunohistochemistry performed on sections of formalin-fixed, paraffin-embedded popliteal and inguinal lymph nodes of one test animal per group provided further evidence of DC-specific transduction, as the ffLuciferase-positive cells had a typical DC morphology (Figure 3B). Interestingly, untargeted adenofection lead also to marginal transduction of DCs. Nonetheless, upon transduction with the α-DC adapter, the histology sample displayed a substantially higher number of positive DCs and the staining for the reporter protein appeared more pronounced in ffLuciferase-positive cells (Figure 3B) than with the untargeted control. There was no evidence of off-targeting to lymphocytes or macrophages, based on the morphology and distribution of positive cells. Based on these results, we could conclude that there is direct and increased *in vivo* targeting of dendritic cells localized in draining lymph nodes.

**Figure 3:**
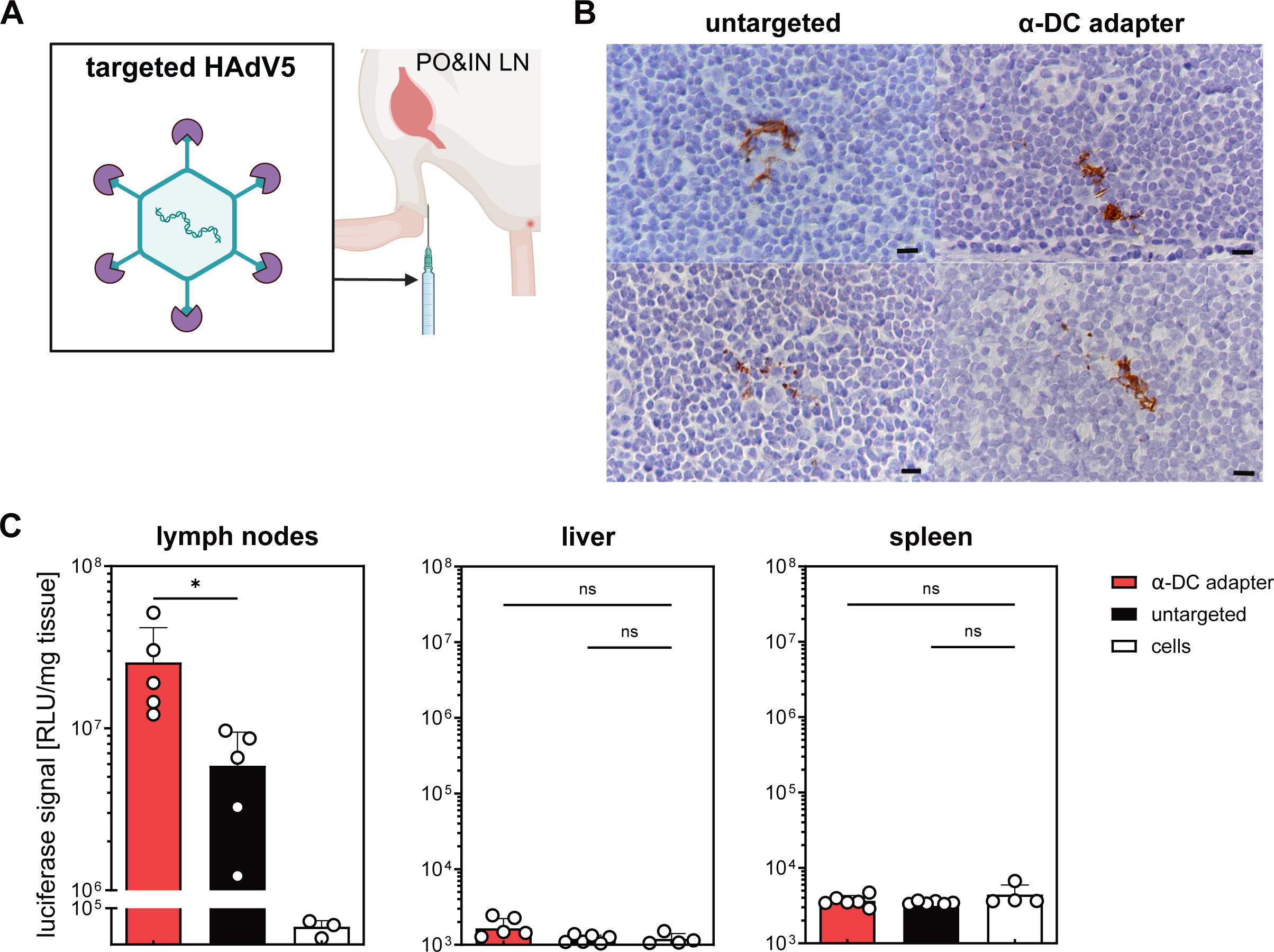
Successful *in vivo* targeting of dendritic cells in draining lymph nodes without off-targeting effects. (A) Schematic representation of *in vivo* biodistribution study. HAdV5^ΔHVR7^ retargeted with α-DC adapter was injected in the hock of the mice (six test animals per group). Two days post injection, popliteal and inguinal lymph nodes, as well as liver and spleen, were isolated and processed for the detection of the expressed reporter protein ffLuciferase. Lymph nodes of one test animal per group were paraffin-fixed and used for IHC. The residual tissues (n=5-6) were lysed to measure luciferase activity. Immunohistochemical staining of lymph nodes for the reporter protein luciferase (of one mouse per group). Two representative lymph node sections are shown for each group. Representative positive staining confirm the transduction specificity to dendritic cells and do not provide any evidence for off-targeting to macrophages and T or B lymphocytes. A higher number of positive DCs and more intense staining for the reporter protein is seen with the α-DC adapter. (C) Luciferase activity in lysed tissue samples (lymph nodes, liver, spleen), normalized by tissue weight. Increased luciferase activity (5-fold) with the α-DC adapter in popliteal and inguinal lymph nodes is seen in comparison to the untargeted HAdV5^ΔHVR7^ control. No off-targeting signal was detected in liver or spleen tissue. Statistics: Each data point represents a single mouse. P-value determined using two-tailed paired t tests or one-way ANOVA with Dunnett’s test for multiple comparisons to tissue background control. *p < 0.05, **p < 0.01, ***p < 0.001, ****p < 0.0001. Bar graphs represent mean ± SD, n = 3-5.

### IL-2v and IL-21 stimulate antigen presentation of HC-HAdV5-transduced dendritic cells in an autocrine manner

As we successfully established DC transduction in draining lymph nodes, we wanted to confirm the therapeutic efficacy in a subsequent tumor vaccination study. One crucial aspect of a successful vaccination is the presentation of an antigen by DCs. In our study, we intended to directly transduce DCs by retargeted adenovirus and deliver the genetic information required for the expression of the model antigen ovalbumin. Generally, antigen processing and presentation entail multiple steps and thus multiple options for delivery. The antigen may be secreted into the host cell’s microenvironment, expressed in the cytosol of the host cell or trafficked to a specific cell compartment. Prior research has established that fusing an antigen with the invariant chain is particularly advantageous for presentation through both MHCI and MHCII of the targeted host APC.^70^ Therefore, we wanted to characterize this approach for OVA peptide presentation upon HAdV5 delivery. We thus transduced DC2.4 cells either with the fusion construct li-OVA1+2 (for details, see Supplementary Information) or with w.t. ovalbumin, which is secreted due to its internal signal sequence that is not cleaved off^71^. Similar to the findings in the literature, the invariant chain fusion accomplished better presentation of the antigen via MHCI compared to secreted w.t. ovalbumin (Figure S7).

To explore the effect of co-stimulatory cytokines, we cloned and produced various high-capacity vectors that contained the encoded antigen li-OVA1+2, and we additionally included an expression cassette for the secretion of a stimulating cytokine (IL-2v or IL-21). The HC-HAdV5 were produced as described previously and purified by two-step CsCl gradient ultracentrifugation to separate HC-HAdV5 vectors from helper virus and producer cell DNA (Figure S8).^19^

We encoded IL-2v or IL-21 for immune stimulation^72-75^ as they alter antigen processing and presentation while potentially enhancing the effect of vaccination (see following section). The same quantity of transducing particles was added to DC2.4 cells. Two days after transduction, we performed pMHCI staining to quantify antigen presentation. Flow cytometric analysis demonstrated that co-expression of IL-2v and IL-21 increased the median fluorescent intensity of MHCI-bound peptide SIINFEKL in the overall population compared to the vector solely encoding the antigen (Figure 4A).

**Figure 4:**
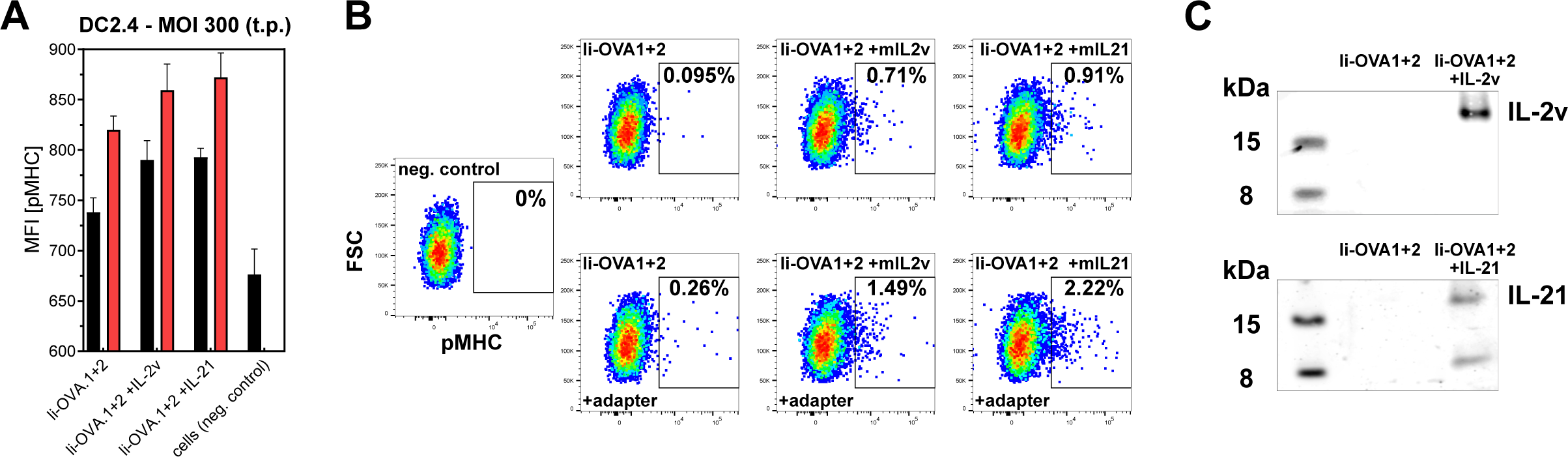
Co-expression of IL-2v and IL-21 stimulate antigen presentation of HC-HAdV5 transduced DC2.4 dendritic cells. Transduction of DC2.4 cells (MOI 300) with HC-HAdV5^ΔHVR7^ encoding the antigen li-OVA.1+2 under the CMV promotor, with and without the co-expression of cytokine IL-2v or IL-21 under the PGK promotor. (A) After two days post transduction, the processed reporter antigen (ovalbumin) was detected as presented antigen peptide (pMHC), using flow cytometry analysis with representative median fluorescent intensity of pMHC staining of the overall population. Black bar: Transduction without adapter (untargeted). Red bar: Transduction with α-DC adapter. t.p.: transducing particles. (B) Flow cytometry analysis of pMHC staining (DC2.4 cells), gating for high antigen-presenting cell population. *Top row*, untargeted; *bottom row*, with α-DC adapter; *negative control*, cells only without vector. (C) Western blot of cell supernatants to confirm secretion of IL-2v (kDa 17) and IL-21 (kDa 15) two days post transduction. No cytokine secretion was detected in the li-OVA1+2 control group (middle lanes). Statistics: Representative data of two independent experiments shown. Bar graphs represent mean ± SD, n = 4

We further separated this population into low and high antigen-presenting cells (Figure 4B). Upon co-expression of IL-2v and IL-21 the population of the high antigen-presenting cells was enriched to 0.91%, while the untargeted il-OVA1+2 control (without co-expression of cytokines) resulted in an enrichment of only 0.095%. By adding the α-DC adapter, we increased the initial transduced population, which resulted in an increase in both the median fluorescent intensity (MFI) of pMHCI staining and the proportion of high antigen-presenting cells (from 0.26% for li-OVA1+2 to 1.49% for li-OVA1+2 with IL-2v or to 2.22% for li-OVA1+2 with IL-21). Consequently, we were able to boost the population of high antigen-presenting cells by 16 to 23-fold by adding the α-DC adapter and co-expressing an appropriate cytokine. The successful expression of the co-encoded cytokines was confirmed by western blot analysis for murine IL-2 or IL-21 staining of the cell supernatants (two days after transduction).

### DC-targeted vaccination with cytokine co-expression leads to efficient suppression of metastases

Having established a DC-targeting adenoviral gene therapy platform that enhances local transgene expression in draining lymph nodes, we hypothesized that DC transduction and direct expression of an antigen in an APC would lead to an increase in vaccination efficacy. Typically, adenoviral vaccination studies are performed using 1^st^ generation adenoviral vectors. However, we were also interested in the performance of high capacity-vectors and their suitability as an antigen delivery tool as they offer much wider possibilities for combinatorial approaches. Our platform aims to express the transgene locally in the draining lymph nodes without off-targeting in liver or spleen. This allows for the local co-expression of potent vaccination adjuvants, owing to the HC-vector’s high coding capacity.

Cytokines serve as strong inducers in immune therapy, however, their potential inflammatory side-effects constrain their therapeutic potential when administered systemically. Controlled local expression at the site of action is therefore particularly important, and it was previously shown that our system can lead to high localized expression and low systemic concentrations.^40^ As we showed evidence of improved antigen presentation by IL-2v and IL-21 secretion (see above), we also wanted to test the beneficial impact towards the overall vaccination process *in vivo*. We hypothesized that transduction of DCs and co-expression of IL-2v and IL-21 results in further stimulation of surrounding immune cells in a paracrine manner. IL-2v is a mutated form of murine interleukin 2 that only binds to IL-2Rβ and IL-2Rγ (but not to IL-2Rα), resulting in suppressed Treg activation and reduced Fas-mediated apoptosis^72,73^. IL-2v was shown to expand NK and CD8^+^ T cell pools^72,73^ with the indication of also having a positive impact on long-term memory T cell formation due to the preferential binding to IL-2Rβ and IL-2Rγ but not to IL-2Rα^76^.

IL-21 has a wide-ranging effect on various immune cells. For instance, it induces the proliferation of cytotoxic CD8^+^ T cells, the differentiation of plasma cells, increased immunoglobulin production in B cells, and maturation and enhanced cytotoxicity of NK cells^74,75^. Additionally, IL-21 has been found to promote the development of memory T cells.^74,75^ Overall, both cytokines may have a positive impact on both tumor regression and long-term memory formation. To test this hypothesis, we cloned and produced HAdV5 and HC-HAdV5 vectors encoding the antigen li-OVA1+2 alone or encoding the antigen in combination with the cytokines IL-2v and IL-21 (Figure S8).

We performed two tumor vaccination studies using the B16-OVA MO4 lung metastasis model in C57BL/6 mice, exploring administration of either low or high vector dosage. After two adenoviral vector administrations with low vector load injections of 1×10^7^ transducing vector particles (vp), or high-load injections of 3×10^8^ transducing vector particles (vp), we examined the lungs at the scheduled end point of 21 days (Figure 5A). We assessed several parameters to robustly determine the extent of metastasis formation and tumor growth in the lung: lung weight, the number of metastatic foci visible beneath the pleura upon macroscopic examination, and determination of the melanin content in a lung cell lysate (Figures 5B and 5D). Furthermore, we quantitatively assessed sections from the lungs of one animal per treatment group via immunohistochemistry by staining for the melanoma marker gp100^77^, followed by morphometric analysis to quantify the proportion of lung tissue occupied by tumor tissue metastases (Figure S9).

**Figure 5:**
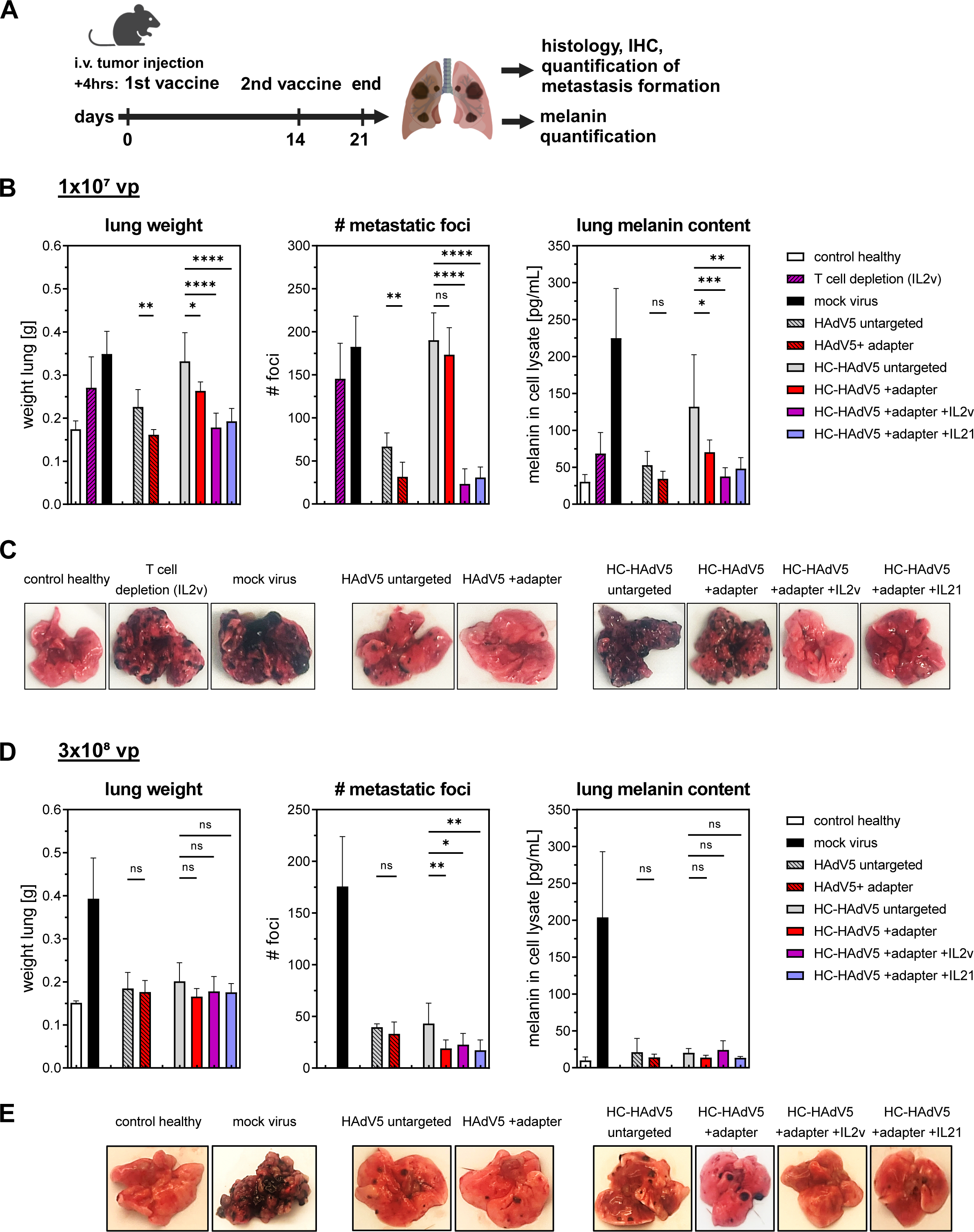
Dendritic-cell targeted adenoviral vectors are more efficient as tumor vaccines. (A) Schematic representation of *in vivo* tumor vaccination efficacy study. After tail vein injection of 4.5×10^5^ B16-OVA MO4 cells at day 0, mice were vaccinated subcutaneously two times (4 h post *i.v.* injection and at day 14). Adenoviral vectors encoded the antigen li-OVA.1+2 under the CMV promotor, with and without co-expressed IL-2v or IL-21 under the PGK promotor. At the end of the experiment at day 21, isolated lungs were examined for tumor treatment. The study was done with low vector load injections of 1×10^7^ transducing vector particles (vp) (B,C), or high-load injections 3×10^8^ vp (D,E). (B,D) Tumor growth analysis of harvested lungs at the end point at day 21. Tumor regression was measured based on lung weight, number of superficial metastatic foci and melanin content of lung lysate. One treatment focused on HAdV5 vaccination (1^st^ gen vector, denoted “HAdV5”), in which one of the two sample groups was targeted (α-DC adapter, denoted “+adapter”), the other was not (“untargeted”). The next treatment focused on HC-HAdV5 vaccination (gutless vectors, denoted “HC-HAdV5”), in which three of the four sample groups were targeted (α-DC adapter), one was not. Two of the targeted HC-HAdV5 vectors also encoded the murine cytokine IL-2v or IL-21 under the PGK promotor (in addition to the encoded antigen li-OVA1+2). As control groups, mice were injected with mock virus (no transgene encoded). To test the effect of CD8+ T cells, these cells were depleted in a treatment with HAdV5 +adapter +IL-2v (“T cell depletion”). Healthy mice without treatment and without tumor burden were taken to assess treatment success. (C,E) Representative visualization of the lungs at the end point measurement. Statistics: P-value determined using two-tailed paired t tests or one-way ANOVA with Dunnett’s test for multiple comparisons to untargeted control. *p < 0.05, **p < 0.01, ***p < 0.001, ****p < 0.0001. Bar graphs represent mean ± SD, n = 5-7.

The results showed pronounced suppression of lung metastases in mice treated with retargeted HAdV5 or HC-HAdV5 (antigen li-OVA.1+2) compared to the untargeted control (Figures 5B-E). In the HAdV5 low vector dosage group, the beneficial retargeting effect of the α-DC adapter led to the presence of only minimal metastatic growth, with only a few small metastatic foci detected (Figures 5B and 5C) (Figure S9F). Morphometric analysis of the lungs indicated that only 0.15% of the lung area displayed gp100 expression (Figure S9A). While the mice treated with the untargeted HAdV5 also exhibited some reduction in tumor burden when compared to the mock control with numerous metastatic foci (Figures 5B and 5C) (Figures S9D and S9E), the treatment effect was significantly less substantial compared to the effect of the targeted HAdV5 group.

For the untargeted group of the HC-HAdV5 in the low dosage study, numerous variably sized metastatic foci were seen (Figures 5B and 5C) (Figures S9B and S9G), and no prominent treatment effect was found compared to the mock control group. In contrast, when the α-DC adapter was added, a significant reduction in lung weight and lung melanin content was observed (Figure 5B). Moreover, the co-expression of mu IL-2v or mu IL-21 achieved substantial further enhancement of the vaccination response for the HC-HAdV5 α-DC adapter groups. By co-expressing those cytokines, hardly any tumor tissue was detectable (Figures 5B and 5C), with only few small subpleural and parenchymal metastatic patches (Figures S9I and S9J).

The growth reduction of the B16-OVA MO4 metastasis model is described to be mainly dependent on cytotoxic CD8^+^ T cells, however, it is also under strong control of NK cells.^78,79^ IHC staining of CD8 and CD161 revealed that NK and CD8^+^ T cells were seen in variable numbers within metastatic foci in all treatment groups. However, for progressed large metastatic foci, both cell types failed to reach the center (IHC data of CD8 and CD161 not shown). IL-2v (or IL-21) can also stimulate NK cells, and to test which cell type is mostly responsible for tumor control in this model, we depleted CD8^+^ T cells prior to administering of DC-retargeted HC-HAdV5 that were co-encoding for IL-2v. The tumor burden was close to that of the mock control for lung weight and metastatic foci, however, less total melanin was measured in the lung lysate (Figure 5B). Immunohistochemistry and morphometric analysis of gp100 expression (Figure S9A,C,D) indicated that 20% of the lung area stained for gp100 in the mock-treated animal, and only 11% in the T cell-depleted animal, suggesting a partial effect by the NK cells alone. With administration of retargeted HC-HAdV5 (co-encoding IL-2v), very limited metastasis formation was seen by histology, and now only 0.7% of the lung area expressed gp100. Staining of the lung for CD8 and CD161 of the T cell-depleted animal confirmed a total absence of CD8-positive T cells, while in several metastases NK cells were detected between neoplastic cells. Therefore, we conclude that in our case the treatment in the cytokine-stimulated groups was indeed mainly based on cytotoxic CD8^+^ T cells, while stimulated NK cells contributed additionally to the treatment success.

The beneficial effects of targeted DC transduction and co-expression of murine IL-2v or IL-21 could also be observed in a high-dosage study, resulting in minimal metastatic tumor growth in the targeted HAdV5 and HC-HAdV5 groups (Figures 5D and 5E). However, due to the higher vector load administered, differences between the treatment groups became less evident. Interestingly, the measured parameters (lung weight, metastatic foci and lung melanin content) of the untargeted HC-HAdV5 group were still above the healthy control for the high dosage (3×10^8^ vp) (Figures 5D and 5E), while for the low dosage study the targeted HC-HAdV5 +cytokine groups displayed only minimal metastatic tumor growth (Figures 5B and 5C). Based on these results we conclude that retargeting to dendritic cells when including the stimulation by secreted cytokines results in better treatment outcomes during tumor vaccination, even at a 30-fold lower dosage than its untargeted vaccination control.

### Co-expression of IL-2v or IL-21 by transduced dendritic cells boosts adaptive response

We observed in our tumor vaccination study a significant suppression of metastasis formation in the treatment groups with vector retargeted to DCs. Particularly, the combination therapy with co-expressing IL-2v and IL-21 in lymphatic tissue resulted in the most efficacious treatment. We measured the level of antigen-specific T cells in the spleen by binding to the MHC I dextramer, since in the B16 melanoma model, cytotoxic T cells are the primary effectors mediating tumor cell killing. Our measurements confirmed a correlation between higher levels of antigen-specific T-cells and improved treatment outcomes.

Our study aimed to investigate whether DC-directed vaccination supports the proliferation of antigen-specific T cells (Figure 6A). The quantified population size of antigen-specific T cells isolated from the spleen indeed supported this hypothesis, since the mean values of this population were found to increase by 50-90% upon DC-targeted transduction, compared to the untargeted control (HAdV5 and HC-HAdV5). This finding corroborated our hypothesis that DC-targeted vaccination promoted the proliferation of antigen-specific T cells. Retargeting of HC-vectors in combination with co-expression of cytokines increased the level of cytotoxic T cells by 330-400%, compared to the untargeted HC-HAdV5 control (Figure 6C). This trend was seen for both the high and low vector load vaccination, confirming our suggested mode of action in both cases. When we isolated splenocytes and stimulated them with peptide antigen, we observed a peak response in IFNγ secretion for the retargeted HC-HAdV5 treatment group with co-expression of IL-2v or IL-21 (Figure 6B). Although expected due to the previously measured increase in cytotoxic T cells, the level of released IFNγ was significantly higher (≥20-fold) than the increase in numbers of specific T cells. This indicated that IL-2v and IL-21 not only promoted CD8^+^ T cell proliferation, but that the resulting T cells were also particularly reactive.

**Figure 6:**
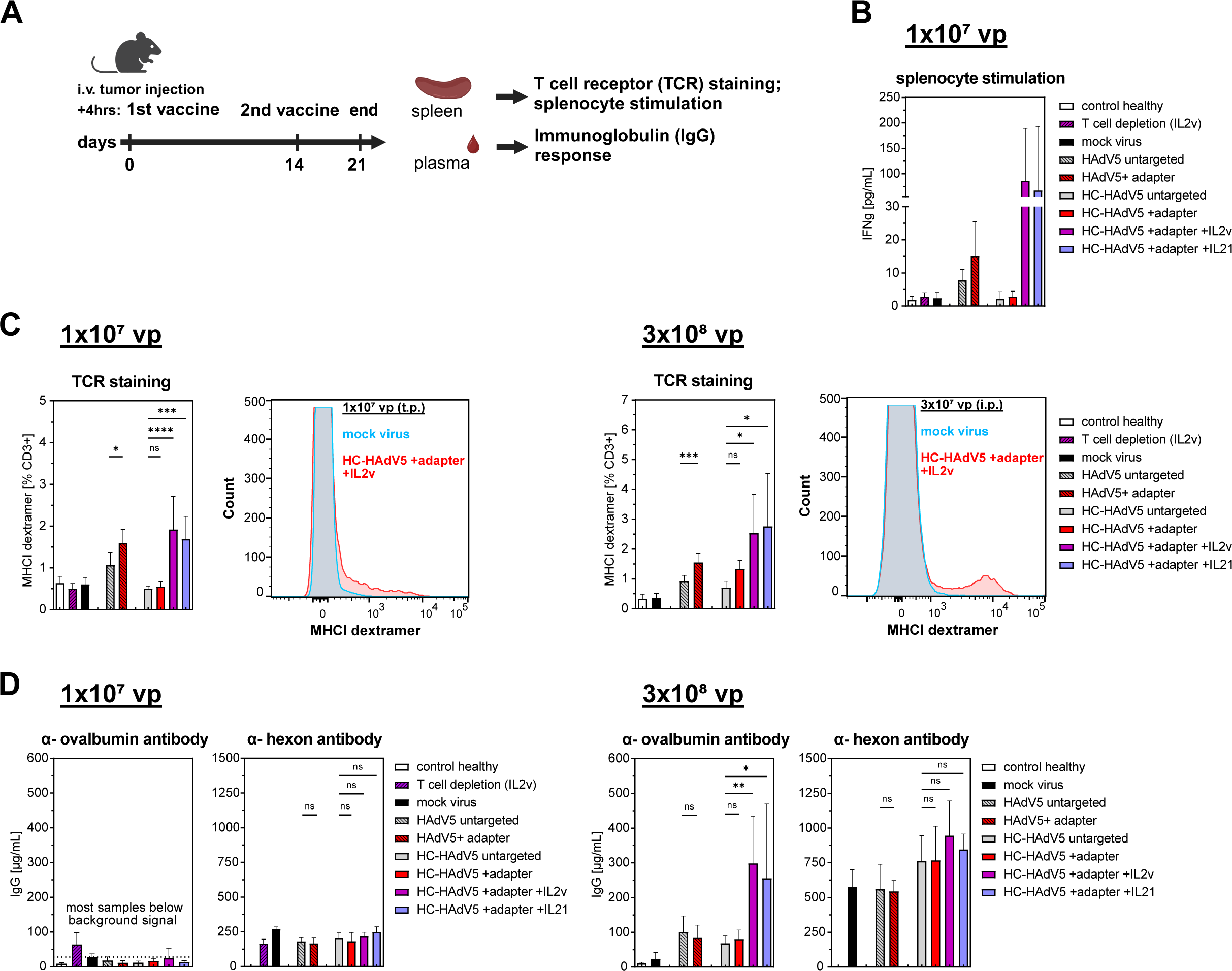
Dendritic cell-targeted adenoviral vectors alter adaptive immune response. (A) Schematic representation of *in vivo* tumor vaccination efficacy study. After tail vein injection of 4.5×10^5^ B16-OVA MO4 cells at day 0, mice were vaccinated subcutaneously two times (4 h post *i.v.* injection and at day 14). Adenoviral vectors encoded the antigen li-OVA.1+2 under the CMV promotor. At the end of the experiment at day 21, isolated spleens and blood plasma were examined for adaptive T and B cell response. (B) Splenocyte stimulation (post low-load treatment of 1×10^7^ vp) with ovalbumin peptide SIINFEKL as CD8^+^ T cell epitope. Detection via ELISA of secreted IFNγ. Groups are explained in Fig. 5B. (C) Flow cytometry analysis. Quantification of cytotoxic T cells in the CD3^+^ population by staining with SIINFEKL MHC-I dextramer, in the low-(1×10^7^ vp) and high-load treatments (3×10^8^ vp). (D) ELISA quantification of anti-ovalbumin IgGs or anti-hexon IgGs in blood plasma. Samples from the low-(1×10^7^ vp) and the high-load treatment study (3×10^8^ vp). Samples for α-ovalbumin IgGs in the low viral load study were below the detection limit. Statistics: P-value determined using two-tailed paired t tests or one-way ANOVA with Dunnett’s test for multiple comparisons to untargeted control. *p < 0.05, **p < 0.01, ***p < 0.001, ****p < 0.0001. Bar graphs represent mean ± SD, n = 5-7.

Besides the T cell response, we additionally explored the B cell response by measuring the titers of antibodies binding to ovalbumin or adenoviral hexon protein (Figure 6D). Interestingly, the adenoviral DC-targeting alone, even at high-load vaccination, had little effect on the immunoglobulin G (IgG) response against the encoded antigen ovalbumin; the titers were on the same level as their untargeted HAdV5 or HC-HAdV5 counterpart. In contrast, the secretion of IL-2v or IL-21 by transduced host cells into the lymphatic microenvironment increased the IgG levels by 3 to 4-fold. Importantly, this increase was limited to the IgG response against the DC-expressed antigen ovalbumin, while the IgG response against the hexon, the major capsid protein, showed no significant difference: the release of IL-2v or IL-21 had no significant effect on the level of anti-hexon IgG, the levels were similar to those in the targeted and untargeted HC-HAdV5 control. In light of repeated vector administrations, a directed and increased antibody response towards the expressed antigen but a missing increase in humoral response towards the vector would be of particular importance to avoid the risk of virus neutralization. In summary, we can conclude that the success of a DC-targeted vaccination can indeed be further improved by the secretion of cytokines into the lymphatic microenvironment, resulting in an increased humoral response directed to the antigen expressed by the host DC.

## Discussion

Human DCs are naturally resistant to infection by human adenovirus C5 due to the lack of CAR and a_v_β_5_ integrin expression. However, transduction of DCs can have very beneficial effects on immune therapy, as we showcased in our study. We established targeted transduction of DCs by human adenovirus C5 specifically in draining lymph nodes and thereby stimulated lymph node residential immune cells in a paracrine manner, exploiting the co-expression of cytokines. This study illustrates a new way of potent combination therapy in a directed tumor vaccination approach, which allowed drastic reduction of adenoviral vector titers for a potent therapeutic effect.

We extended our adenoviral gene therapy platform to murine and human DCs by designing a retargeting adapter that uses DC-SIGN (CD209) and CD11c as novel entry receptors. The dual-adapter system, in which two retargeting modules redirect HAdV5 to specific surface markers on primary human and murine DCs, led to an increased transgene expression by 5 to 17-fold for at least 10 days (Figures 2F and 2H). A dual-adapter system offers the advantage to allow better transduction of cell types that feature medium or low expression levels of the specific cell markers, such as DC-SIGN or CD11c. We found a significantly enhanced α-DC adapter-mediated transduction efficiency by fusion of a CD11c-binding peptide to the DC-SIGN retargeting domain (GP1_LASV_). We did not further investigate the detailed synergistic transduction effects but hypothesize that two retargeting modules led to avidity effects. In addition, receptor clustering could be induced on the cell surface by the trimeric adapter, as each of the two retargeting modules could lead to receptor cross-linking.^80^

The retargeting module to DC-SIGN consists of GP1_LASV_ and includes a point mutation (M192R).^60^ The LASV, from the family of the arenaviruses, was previously described to infect human monocytes and DCs by binding via GP1 to its primary entry receptor DC-SIGN (and dystroglycan).^43^ However, GP1 of LASV also binds to LAMP1 in late endosomes, as part of the initiation of pore formation of arenaviruses. To utilize GP1_LASV_ as a retargeting module for HAdV5 transduction, we identified the benefits of eliminating the GP1 binding to LAMP1 (Figure 1B). During a natural infection of lung epithelial cells, HAdV5 particles very rapidly traffic to the nucleus and only minor amounts can be co-localized in early or late endosomes 2 hours post-infection.^81^ Natural viral trafficking of HAdV5 in DCs has not been investigated further due to the lack of natural infection, but might be influenced by the specific endosomal pH milieu in DCs compared to other cell types.^82^ We hypothesize that especially at higher MOIs the endosomal escape of HAdV5 in early endosomes is hindered by rebinding of the retargeting adapter to LAMP1, resulting in the potential degradation in endolysosomes. Therefore, the elimination of LAMP1 binding was a design goal, and the observed improvements are consistent with our hypothesis.

The concept of retargeting HAdV5 is a well-studied field.^37,40-42,83-85^ Nevertheless, with the dual targeting approach and the optimal set of retargeting moieties, we may have found an improved approach to target HAdV5 specifically to DCs, allowing directed transduction even at low vector loads. Besides targeting of HAdV5 vectors to a defined cell type we also reduced *in vivo* off-targeting effects. We deleted the hexon HVR7 epitope which ablates binding to the blood coagulation factor X (FX), and used a dual retargeting adapter which sterically covers the CAR epitope on the fiber knob. Thus, we achieved preferential transduction of DCs in murine lymph nodes while not transducing other cell types like B or T cells, which both outnumber our target cell by far in lymph nodes (15% B cells and 78% T cells vs 0.5% DCs) (Figure 3B).

In turn, the many challenges of an *in vivo* targeting of DCs have so far not yet been fully overcome with other approaches. For example, DC-SIGN,^86,87^ DEC-205^88^ and CD40^89-91^ were previously used in alternative adenoviral targeting systems; however, targeting of those markers was reported to be successful only at very high vector loads (≥MOI 300) or not even confirmed in a direct *in vivo* vaccination. Our own experiments revealed that the tested constructs targeting DEC-205 or CD40 were not efficient in inducing HAdV5 transduction (Figure S2). Although DEC205 remains a promising surface marker, further investigations are required to specifically select a binder for this receptor to mediate transduction, as not all binders do this — as previously shown for hFAP.^39^ The receptor CD40, on the other hand, displays features that do not make it particularly suitable as a potential receptor for DC targeting. It is only expressed at low levels on immature DCs, and targeting of this receptor would be accompanied with off-targeting to CD40-positive B cells, especially in lymph nodes where B cells are far more numerous than DCs. Several HAdV5 vaccinations which CD40 targeting were subsequently performed *ex situ* instead of *in vivo*.^91,92^

In contrast to knob-directed retargeting strategies, lactoferrin offers a capsid-mediated transduction enhancement to DCs: Lactoferrin in high concentrations is a popular reagent to transduce DCs *in vitro*, as it covers the particle via electrostatic interactions to the negatively charged hexon and binds to DC-SIGN in a bispecific manner.^36^ Nevertheless, our α-DC adapter outcompeted lactoferrin at the tested MOIs in direct comparison experiments. Furthermore, the HAdV5-lactoferrin complex mediates activation of APCs through TRL4 stimulation, resulting in inflammatory responses with release of TNF-α and IL-1β^26^ and therefore limits therapeutic options. Additionally, lactoferrin does not uniquely bind to DC-SIGN on APCs but also interacts with other receptors (e.g. CD91, CD14, Intelectin-1) on multiple cell types *in vivo*^93^.

Various DC-targeted gene therapy strategies for tumor vaccination have previously been pursued by others.^94,95^ While AAVs and lipid-nanoparticles feature a limited packaging capacity, lentiviral systems can integrate into the patient’s genome.^9^ On the other hand, there is a beneficial retargeting effect that is mediated by GP1_LASV_. It is therefore worth considering whether LASV can be used as a viral delivery system for the transduction of DCs, potentially competing with HAdV5 for DC gene therapy purposes. While LASV can infect and replicate in DCs, infected DCs lack the ability to activate T cells due to the immunosuppressive nature of the LASV nucleoprotein.^96^ A LASV-based vector therefore would need much further engineering to make it as safe as the clinically proven non-replicating 1^st^ generation HAdV5. Currently, other vectors belonging to the family of arenaviruses are being developed for vaccination approaches (e.g. lymphocytic choriomeningitis virus). However, the replacement of the encoded glycoprotein on the S segment offers only very limited capacity for an antigen with a maximum 1-2 kbp size, and therefore multiple transgenes need to be encoded on multiple vectors.^97,98^ In contrast, an adenoviral vaccination system, specifically HC-HAdV5, has the unique advantage of being non-integrative and non-replicative while providing a packaging capacity of up to 37 kbp.

Our work is primarily focused on the gene delivery of an intracellularly expressed antigen to DCs. Nuclear delivery of the DNA encoding the antigen, followed by antigen expression, direct processing and antigen presentation by an APC represents a potent way of cytotoxic T cell proliferation. The directed antigen trafficking in MHC I and MHC II loading compartments by using fusions to invariant chain is state of the art and broadly applied in the processing of multimer-neoantigens.^99^ In contrast, the endocytosis of an extracellular antigen with subsequent processing is less efficient, as only a minority of antigen molecules are finally presented as epitopes.^100^ Our tumor efficacy study displayed the advantages of targeting DCs and *in situ* expression of a desired antigen in draining lymph nodes.

We particularly showed evidence of increased antigen presentation by autocrine stimulation of co-expressed IL-2v or IL-21 (Figure 4). While the application of IL-2 as stimulator for DC antigen presentation is undisputed, the impact of IL-21 in immune therapy has been controversially discussed. It had been reported that exocrine stimulation of murine DCs with IL-21 leads to decreased antigen presentation, resulting in dysfunctional T cell priming.^101^ However, the role of IL-21 becomes more complex as murine GMCSF-DCs (to which the DC2.4 cells belongs)^102^ respond differently to exocrine IL-21, and this effect is also reported differently for monocyte-derived human DCs.^103^ In line with our results, two other groups highlighted in DC-targeted vaccination studies that autocrine IL-21 signaling led to an increase in the antigen presenting cDC1s population,^104^ suggesting that targeted DCs did not fail to stimulate CD8^+^ T cells.^105^ Furthermore, the local secretion of IL-2v and IL-21 in our study led to distinct enhancement of cytotoxic T cell proliferation and activity (Figure 6). Consequently, the tumor burden was significantly reduced with a low adenoviral vector load of only 1×10^7^ vp (transducing vector particles), whereas other tumor vaccinations are typically administered within a range of 1×10^8^ to 1×10^9^ vp.^88,106^ We hypothesize that the observed cytokine booster effect is most effective in a lymphatic environment. As the targeted DCs migrate to draining lymph nodes and interact locally with T cells, B cells and NK cells while secreting the encoded cytokines, DCs become a potent host cell for paracrine delivery of immune stimulants. Our lymph node-targeting strategy has multiple potential applications as the advantageous effect of immune stimulants (e.g. checkpoint-blockade release) in draining lymph nodes was previously described.^107^

In addition to their role in the T cell response, IL-2v and IL-21 also play essential roles in B cell expansion and immunoglobulin production. Interestingly, the IgG levels with specificity against the encoded intracellular antigen (anti-ovalbumin IgG) increased by factor 3-4 in the treatment groups with IL-2v or IL-21, while anti-AdV5 hexon IgG levels did not increase upon co-expression of the described cytokines. The adaptive antibody response is complex, as many cell types are involved (B cells, T helper cells, DCs, macrophages), nevertheless, this finding supports the idea of directing antigen delivery to DCs with paracrine cytokine stimulation to T helper cells and B cells. As the encoded antigen was expressed under a strong promotor and presented over a long time period, we presumed that it out-competed AdV5 epitopes of the delivered particle in a transduced DC. In contrast, unsuccessful transducing vector particles (i.e. particles not leading to expression of antigen and co-stimulatory cytokines) would be degraded by macrophages and only evoke anti-viral IgG responses. Anti-viral immunity, as it is introduced by repeated vaccinations, can become a substantial drawback for adenoviral vectors. We are well aware that capsid covering can be required for repeated booster vaccination, and we have therefore previously developed and characterized a hexon-covering protein shield.^41^ While this aspect was not in the scope of our DC-targeting study, we assume that by shielding a booster vaccine, the levels of anti-hexon IgGs can decrease further and overall vaccination might be even more successful due to the decreased neutralization effect. Currently, we could demonstrate that a shielded and targeted HAdV5 could still efficiently transduce murine and human DCs (Figure 2).

## Limitation of study

In our study we have worked both with 1^st^ generation adenoviral vectors (HAdV5) and gutless or high-capacity vectors (HC-HAdV5). Both vector generations were able to evoke an adaptive anti-tumor response (Figures 5 and 6). We refrain from making a direct comparison between HAdV5 and HC-HAdV5, and in our tumor study with high vector load, only a marginal difference was found between the vectors of different generations. In a direct comparison, several production batches would have to be tested and detailed analyses of the evoked innate immune response performed in order to reliably demonstrate and claim potential differences between HAdV5 and HC-HAdV5 as a vaccination vector. This was not the scope of our study. Clearly, the HC-HAdV5 is well suited as a vector for vaccination and the high transgene capacity is particularly attractive for combination treatment as demonstrated by our work.

## Materials and Methods

### Design, expression and purification of retargeting adapters

Retargeting modules (scFv, peptides or viral glycoproteins (GPs)) binding to CD40, CD11c, CD206, DEC205 (CD205) and DC-SIGN (CD209) were cloned as mono-adapter or dual-adapter format into the expression plasmid pcDNA3.1, analogous to previously described constructs^37,42^. The N-terminal retargeting module is preceded by an HSA leader peptide and followed by a flexible linker, followed by the knob-binding DARPin 1D3nc and the trimerizing domain SHP. Linkers between the two retargeting modules in dual targeting modules and between retargeting module and knob-binding DARPin were in the range of (GGGGS)_4_ to (GGGGS)_6_ (Figure S3B).

From published full-length antibodies, retargeting modules were engineered in the scFv-format (Nterm-VL-(GGGGS)_3_-VH-Cterm). From the published LASV GP sequence (UniProt P08669), either residues 59-237 (GP1) or 59-424 (GP1/GP2), respectively, were used as retargeting modules, binding to DC-SIGN. The introduced point mutation (M192R) blocks binding of LASV GP1 to LAMP1^60^. iCAM1 (UniProt P13597) 407-421 (loop4) was used as retargeting module binding to CD11c^62^.

The adapters were expressed in CHO-S cells via PEI transfection, using 3×10^6^ cells/mL and 5 mg/L PEI mixed with 2.5 mg/L DNA. Cells were incubated at 31 °C for 6 days and 180 rpm. Secreted protein of scFv-containing adapters was directly purified by protein L affinity column from the cell-free supernatant. After 10 column volumes of wash (PBS, pH 7.4), a one-step gradient elution was achieved with elution buffer (0.1 M citric acid, Merck, pH 2.5). Resulting fractions were neutralized immediately with 1 M Tris (Serva, pH 9.0), followed by SEC polishing (see below). In contrast, for secreted protein of non-scFv containing adapters, the supernatant was dialyzed against 20 mM Tris-HCl containing 10 mM NaCl (pH 7.2), using dialysis tubings with a molecular weight cut-off (MWCO) of 12–14 kDa at 4 °C. During 24 h, the buffer was exchanged three times. The dialyzed supernatant was loaded on a Mono Q HR 16/10 column (20 ml CV). After 6 CV wash (20 mM Tris-HCl with 10 mM NaCl), elution was carried out over 20 CV for 0-25% elution buffer (elution buffer: 20 mM Tris-HCl with 1 M NaCl), followed by 100% wash. Product-containing fractions were determined by SDS-PAGE (silver staining). Final polishing of pooled fractions was achieved by SEC on a Superdex 200 with on-column buffer exchange to endotoxin-free PBS (Merck Millipore) and then snap-frozen in liquid nitrogen and stored at -20 °C until use.

### Cell lines and primary cells

The following cell lines were used: murine DC-cell line DC2.4 (Merck SCC142, cultured as recommended by Merck), murine DC-cell line MutuDC^63^ (gift by the Acha-Orbea group, University of Geneva), murine melanoma cell line B16-OVA MO4 (Merck SCC420, cultured as recommended by Merck), human lung cell line A549 (ATCC CCL-185), human embryonic kidney cell line HEK293 (ATCC CRL-1573), 116^108^ (gift by Philip Ng).

Primary mouse cells from the femoral bone marrow of C57BL/6J mice were cultured and differentiated to murine BMDCs.^109^ Human peripheral blood CD14^+^ monocytes cryopreserved (Zenbio SER-CD14-F) were cultured and differentiated to Mo-DC via CellXVivo Human Monocyte-derived DC Differentiation Kit (Bio-Techne CDK004).

For receptor staining by flow cytometry, the following antibodies were used: anti-muCD11c (BioLegend clone N418), anti-hCD11c (BioLegend clone Bu15), anti-muDC-SIGN (BioLegend clone MMD3), anti-hDC-SIGN (BioLegend clone 9E9A8), anti-muCD206 (BioLegend clone C068C2), anti-muCD40 (BioLegend clone 3/23), anti-muDEC205 (BioLegend clone NLDC-145), anti-muH-2Kb (BioLegend clone AF6-88.5), anti-SIINFEKL bound to muH-2Kb (BioLegend clone 25-D1.16)

### Plasmid constructions, payload design and virus production

The HAdV5^HVR7^ vectors (“1^st^ generation” vectors) were obtained by cloning the gene of interest into the pAdEasy-1 plasmid system from Agilent Technologies. The gene of interest is placed under the CMV promotor and followed by a SV40 terminator. Virus particles were produced in HEK293 cells and purified by a CsCl gradient. For the mock HAdV5^HVR7^ vector, a gene of interest is missing between the CMV promotor and the SV40 terminator, resulting in HAdV5^HVR7^ virus particles unable to produce a payload.

The HC-HAdV5^HVR7^ vectors (“gutless” or “high capacity” vectors) were obtained by cloning the gene of interest into the pUniversal plasmids and further combined with the pC4HSU plasmid by Gibson assembly.^19^ HC-HAdV5^HVR7^ was amplified with the helper virus pmCherryHVR7 containing four mutations in the hypervariable loop 7 of the hexon (I421G, T423N, E424S and L426Y). Virus particles were produced in the cell line 116 and purified by a CsCl gradient.^19^

For the design of the antigen, the invariant chain was fused to chicken ovalbumin amino acids 242-353 (“li-OVA.1+2”) and cloned as a gene cassette under the CMV promotor and followed by the BGH terminator. Alternatively, full length ovalbumin (GenBank: SERPINB14) was cloned as gene cassette under the CMV promotor and followed by the BGH terminator. For two pC4HSU constructs, an additional expression cassette with an encoded cytokine was placed downstream of the li-OVA.1+2 expression cassette. These cytokine genes (muIL-2v or muIL-21) included their native signal sequence and were placed under the PGK promotor and followed by a human β-globin terminator. This mutated version of muIL-2 prevents binding to IL-2Rα and does not show preferential activation of Treg cells.^72,73^ The murine IL-21 gene sequence was translated from its GenBank entry (GenBank: AF254070.1) and then the payload genes were codon-optimized for mouse expression and synthesized by Twist Bioscience.

### Genomic and transducing vector particle determination

Transducing adenoviral vector particles were determined by a transducibility assay on A549 cells as described previously.^19^ HC-HAdV5 genomes from isolated nuclei were quantified by qPCR with specific primers binding in the encapsulation signal-region (5′-GACAATTTTCGCGCGGTTTTAGG-3′), (5′-CACTTCCTCTTATTCAGTTTTCCCGC-3′) and a specific double-quenched probe (5′-TGTAGTAAATTTGGGCGTAACCGAGTAAGATTTGGCC-3′). HAdV5 or HV-impurities in the HC-HAdV5 sample (isolated nuclei) were quantified by qPCR with specific primers binding to the hexon encoding region (5′-GAATAACAAGTTTAGAAACCCCACGGTGG-3′, 5′-GTTTGACCTTCGACACCTATTTGAATACCC-3′) and a specific double-quenched probe (5′-TGACATCCGCGGCGTGCTGGACAGG-3′). All reactions were performed using the PrimeTime gene expression master mix (Integrated DNA Technologies, 105571) and the reaction signals were normalized to the passive dye rhodamine-X (ROX).

Dynamic light scattering (DLS) (Wyatt DynaPro III plate reader) confirmed a ratio of approximately 1 transducing vector particle per 2 viral particles (measurements done by external collaborator Vector BioPharma).

### *In vitro* transduction experiments

For screening and characterization of adapter proteins, 2×10^5^ cells/well (DC2.4, MutuDC or primary cells) were seeded in a 96-well format 6 hours prior to the experiment. Reporter virus was incubated with trimeric adapter protein in a ratio of 1 transducing particle to 600- to 1200-fold adapter protein. Virus and adapter protein were incubated for 2 hours on ice and then diluted with PBS (pH 7.4) and added to the cells at the corresponding MOI (multiplicity of infection). After total incubation for 42 hours on cells, cells were washed with PBS. The following reporter viruses were used in this assay: HAdV5^HVR7^ encoding tdTomato under the CMV promotor and HAdV5^HVR7^ encoding ffLuciferase-IRES-GFP under the CMV promotor. In case of transduction with the encoded reporter protein ffLuciferase, passive lysis buffer (Promega, E194A) was added. After 15 min of incubation at RT, ffLuciferase activity in the lysate was determined by a luciferase assay (Promega, E1500) according to the manufacturer’s instructions. In case of transduction with the encoded reporter protein tdTomato, cells were detached from the plate with trypsin or Accutase and resuspended in PBS+1%BSA+0.1%Na-azide. The percentage of fluorescence-positive cells was directly measured by flow cytometry (FACSymphony 5L).

For uptake blocking, anti-muCD11c antibodies (BioLegend clone N418) and anti-muDC-SIGN (BioLegend clone MMD3) were added in a final concentration of 5 µg/mL. As positive uptake control, human lactoferrin was preincubated at a concentration of 1 µM with the virus sample for 2 hours on ice, before dilution and addition to the cells. For blocking the natural uptake that occurs by binding of CAR to the HAdV5 knob, a non-binding DARPin adapter (E2_5-1D3nc_SHP) was used, which covers the CAR epitope on the fiber knob.

Fold changes in transduction efficiency were calculated based on the following formula: [signal measurement – signal cell background]/[signal untargeted control – signal cell background].

### *In vivo* biodistribution and tumor vaccination study (B16-OVA MO4)

All animal experiments were performed in accordance with the Swiss animal protection law and with approval of the Cantonal Veterinary Office (Zurich, Switzerland), license number ZH084/2022. In all experiments C57BL/6J mice (Charles River, 8- to 9-weeks old) were housed in the BSL2 facility of LASC (Laboratory of Animal Services Center, University of Zurich, Switzerland). Prior to each experiment, an acclimatization period of 1 week took place. CO_2_ euthanasia was performed at the end of the experiment.

Six C57BL/6J mice per group (Charles River, 8- to 9-weeks old) were injected subcutaneously into the right hock with HAdV5^HVR7^, encoding ffLuciferase-IRES-GFP under the CMV promotor, that has been previously incubated for 1 hour at 4 °C either with the retargeting anti-DC adapter (“retargeted”) or without additives (“untargeted”). A total of HAdV5 of 3×10^8^ transducing viral particles in 50 µL was injected. After 2 days, mice were euthanized, and liver, spleen as well as popliteal and inguinal lymph nodes were dissected and snap-frozen (lymph nodes of one test animal per group were paraffin-fixed and used for IHC). Frozen tissue was homogenized at 4 °C (Precellys Evolution homogenizer), followed by tissue lysing by adding passive lysis buffer (Promega, E194A) to a defined amount of tissue homogenate. After 15 min of incubation at room temperature, ffLuciferase activity was determined from the lysate’s supernatant by a luciferase assay (Promega, E1500) according to the manufacturer’s instructions. Normalization was done by a BCA assay (protein content of liquid tissue supernatant) (ThermoScientific REF23252).

Five to seven C57BL/6J mice per group (Charles River, 8- to 9-weeks old) were injected intravenously (*i.v.*) (lateral tail vein) with 4.5×10^5^ B16-OVA MO4 mouse melanoma cells suspended in RPMI medium. Prior to injection, the cells had been co-incubated for 2 days with 10 ng/ml mIFN-γ to achieve better lung colonization.^110^ Four hours post-*i.v.* tumor cell injection, a first vaccination was done by subcutaneous injection (*s.c.*) of HAdV5^HVR7^ or HC-HAdV5^HVR7^ into the right hock. A total of 3×10^8^ transducing viral particles, or 1×10^7^ transducing viral particles, respectively, in 50 µL PBS were injected — targeted or untargeted virus samples coding for the indicated payloads (li-OVA1+2 with or without mIL2v or mIL21). A 2^nd^ dose of vaccination was given on day 14 *s.c.* into the right hock, with a total of 3×10^8^ transducing particles, or 1×10^7^ transducing particles, respectively, in 50 µL PBS injected according to the respective treatment group. CD8^+^ T cell depletion was done for one treatment group (targeted HC-HAdV5^HVR7^, encoding for li-OVA.1+2 + muIL2v) with 0.2 mg mCD8-mIgG2 (InvivoGen 10354-42-01) via intraperitoneal injection per mouse and application. CD8^+^ T cell depletion was done on three consecutive days, before, on and after the day of vaccination, at both vaccination time points. On day 21 post tumor injection, mice were euthanized. Blood samples were taken post-mortem through heart puncture into the right ventricle, centrifuged and the blood plasma stored at -20 °C. Lungs and spleen were dissected. The lungs were weighed, and the visible subpleural melanoma foci were counted. One representative lung per group was fixed in 10% buffered formalin for 48 hours and then stored in 70% ethanol until processing for histology and immunohistology. The remaining lungs and spleens were stored at 4 °C until further processing.

### Tissue preparation from tumor vaccination study

Of homogenized and single cell spleen samples (treated with ACK buffer), 1.2×10^6^ splenocytes per sample and staining were used for the determination of T cell subpopulations by flow cytometry. The following stains were applied: Live/DEAD fixable blue stain kit (Invitrogen REF L34962), MHC dextramer H-2 Kb SIINFEKL-APC (Immudex JD2163-APC), mCD3-AF700 (clone 17A2; BioLegend 100215), mCD3-PE (clone 17A2; BioLegend 100206), mCD4-FITC (clone RM4-5; BioLegend 100509), mCD8a-PE (clone KT15; Santa Cruz Biotechnology sc-53473). The increase of cytotoxic T cell population in % was calculated based on the following formula: ([signal measurement – signal healthy control]/[signal untargeted control – signal healthy control]) × 100.

For the splenocyte stimulation assay, 4×10^5^ splenocytes per sample were incubated (in duplicates) in RPMI medium (+penicillin-streptomycin, +10% FCS) in a 24 well plate at 37 °C for 3 days. T cell stimulation by addition of 5 µM OVA 257-264 peptide (InvivoGen vac-sin) was measured by ELISA of secreted mIFN-γ in cell supernatants (Invitrogen REF 88-8314-22).

From mouse blood samples, plasma was diluted in PBS pH 7.4 and α-ovalbumin and α-AdV5 hexon IgG titers were measured by standard ELISA. 5 ng/µl ovalbumin or 5 ng/µL AdV5 hexon was coated overnight at 4 °C. Blocking was done by casein blocking buffer (Sigma-Aldrich B6429), 0.05% Tween-20 in PBS (pH 7.4) for 1 hour. Sample incubation was done for 1 hour. The detection antibody was anti-mouse IgG (H+L)-HRP (ThermoScientific 31438) and the detection was done with tetramethylbenzidine (TMB) substrate (ThermoScientific 34021). For IgG level quantifications, the reference antibodies mouse anti-ovalbumin antibody clone TOSGAA1 (BioLegend 520401) or mouse α-adenovirus hexon antibody (BioRad 0400-0063) were used as standard proteins.

For the quantification of tumor burden in the lung, the melanin content was measured. The melanin content of homogenized and single-cell lung samples was measured according to the Melanin Assay Kit (Fluorometric) (Abnova REFKA6030).

### Histological and immunohistochemical examinations and morphometric analysis

The formalin-fixed lungs and the lymph nodes were trimmed and paraffin wax-embedded. Consecutive sections (3-4 µm) were prepared and stained with hematoxylin-eosin (HE) for histological examination, or subjected to immunohistochemical staining. For immunohistochemistry (IHC), the horseradish peroxidase (HRP) method was applied as detailed below; all stains were carried out in an autostainer (Dako). The lymph node sections were stained for the expression of firefly luciferase,^111^ whereas the lungs were stained for the expression of the melanoma marker gp100, and to identify CD8-positive T cells and NK cells.

Briefly, sections were deparaffinized, dehydrated and subjected to antigen retrieval (20 min incubation at 98 °C in citrate buffer, pH 6 (Agilent Dako) for CD8, and in EDTA buffer, pH 9 for luciferase and NK1.1/CD161; 10 min incubation at room temperature (RT) in Fast Enzyme (Zytomed) for gp100. After incubation with the primary antibodies (mouse anti-firefly luciferase, clone Luci17; ab16466, Abcam; rabbit monoclonal anti-mouse CD8, 989415, Cell Signaling Technology; rabbit anti-NK1.1/CD161, clone E6Y9G, 39197S, Cell Signaling Technology, for 1 h at RT; rabbit anti-gp100, ab137078, Abcam, overnight at 4 °C) and blocking of endogenous peroxidase (peroxidase block; Dako) for 10 min at RT, sections were incubated for 30 min at RT with the detection system (Envision+ System HRP Mouse and Rabbit, respectively; Agilent Dako), followed by incubation with diaminobenzidine as chromogen and counterstaining with hematoxylin.

All histological and immunohistochemical specimens were examined by a veterinary pathologist (AK) who was blinded to the treatment of the animals.

Lung sections stained for gp100 were subjected to a morphometric analysis. The immuno-stained sections were scanned (NanoZoomer-XR C12000; Hamamatsu) and analyzed using the software Visiopharm (version 2023.09.3.15043 x64) to quantify the area of gp100 expression in relation to the total area (area occupied by lung tissue) in the sections. This was done to compare the extent of gp100 expression as an equivalent of metastasis formation in the lungs between the different animals. After manual outlining of the lung samples of interest (ROIs), a first APP (Analysis Protocol Package) was applied that refined the manual annotation. For this purpose, a Decision Forest method was used and the software was trained to detect the lung tissue in the section (total area), and to exclude the optically empty spaces at the level of the blood vessels, airways, and alveoli, as well as the surrounding background. Subsequently, a second APP based on a threshold method was used to detect gp100 expression (as brown DAB precipitate) within the ROI. The gp100 expression, as well as the total area occupied by lung tissue for each animal, were quantified by a third APP. To calculate the percentage of immuno-stained area for each animal, the results were imported into Excel (Microsoft Office Professional Plus 2019; Microsoft, Redmond, USA) and the following formula was applied: [immuno-stained area (µm^2^)]/[total area (µm^2^)]) × 100.

## Acknowledgments

We want to thank Dr. Petra Seebeck and Dr. Felix Gantenbein (Zurich Integrative Rodent Physiology (ZIRP) team) for supporting the animal license application. We would like to thank the University of Zurich Cytometry Facility and the Laboratory of Animal Services Center as well as the laboratory technician of the Histology Laboratory, Institute of Veterinary Pathology, Vetsuisse Faculty, for excellent technical support. We would like to thank Simon De Neck for his support towards the morphometric analysis. We would like to thank Dr. Dominik Brücher for the conceptualization of the high-capacity vectors, providing material and initial support of the project. We would like to thank Prof. Hans Acha-Orbea for providing cell lines. We are grateful to Prof. Urs Greber, Prof. Christian Münz and PD Dr. Dr. Angelika Riemer for critical discussions and support during the project. This project was funded by the Schweizerischer Nationalfonds Sinergia grant CRSII5_170929 and Cancer Research Institute grant F-41105-35-01 (both to A.P.). The AI writing tool DeepL Write supported the text correction process. Graphical abstract and certain figures were created with BioRender.com

## Author contribution

F.W. and A.P. conceptualized the project; F.W. cloned and purified all proteins, generated and produced the viral vectors, set up methodology and performed all experiments (if not stated otherwise), analyzed and visualized data and wrote the original manuscript; J.K and P.C.F supported the methodology related to viral vector cloning and adapter production, supported *in vivo* sample preparation and edited the manuscript; F.G. applied together with F.W. for an animal license at the Cantonal Veterinary Office, co-designed and performed with F.W. the *in vivo* studies, supported *in vivo* sample preparation and edited the manuscript; A.K. performed, validated and analyzed all histological and immunohistological related data and edited the manuscript; A.P. supervised the project, acquired funding and edited the manuscript.

