## Supplementary Figures for "Dendritic cell targeting in lymph nodes with engineered modular adapters improves HAdV5 and HC-HAdV5 tumor vaccination by co-secretion of IL-2v and IL-21"

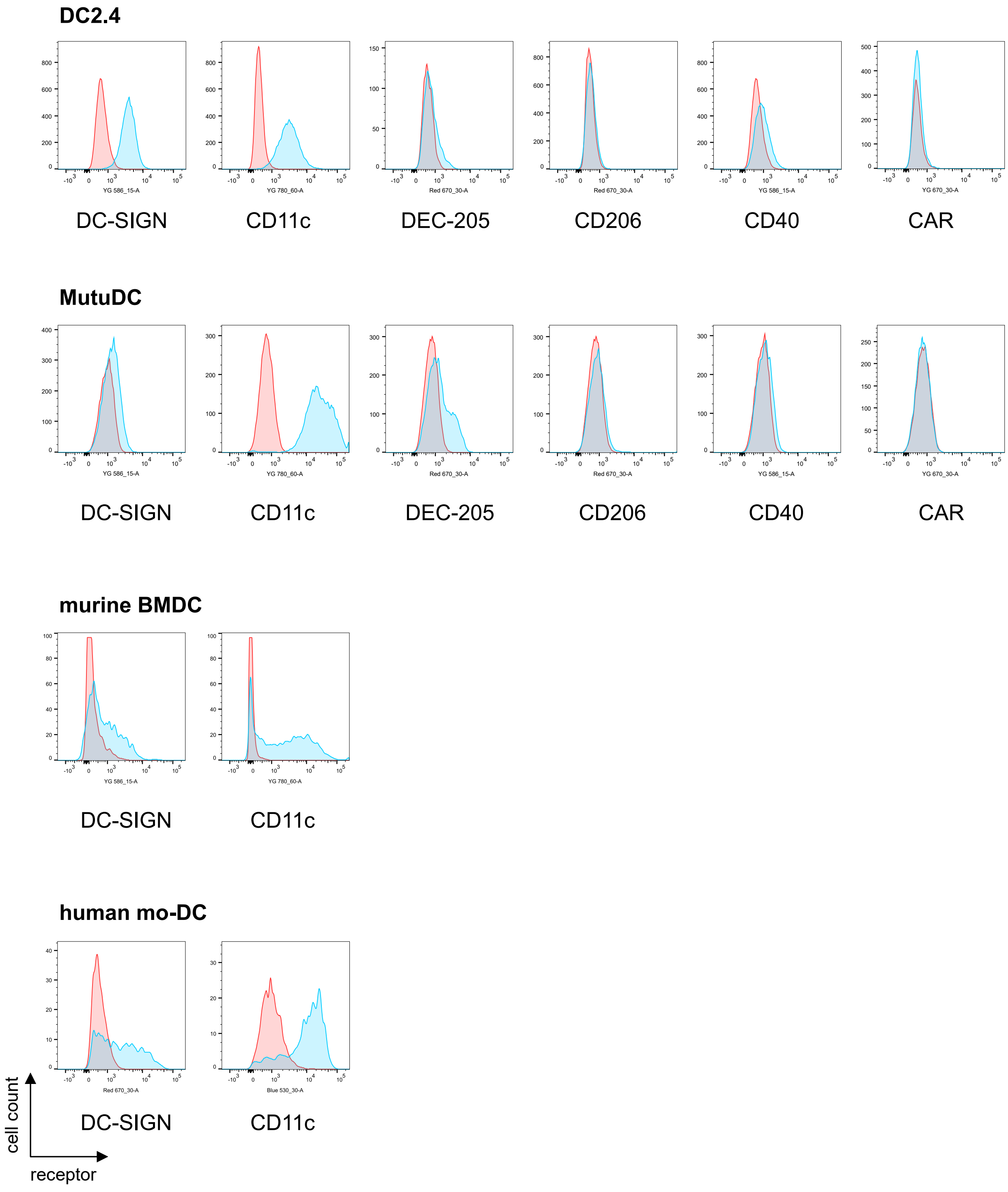

**Figure S1: Histogram of surface receptor staining for DC cell lines and primary cells by flow cytometry**

Flow cytometry analysis of surface marker expression of DC2.4 and MutuDC cell lines, as well as murine bone marrow-derived DCs and human monocyte-derived DCs. Red = unstained control; blue = staining of specific surface marker (antibodies are listed in Material and Methods)

A

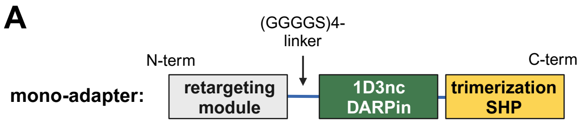

D

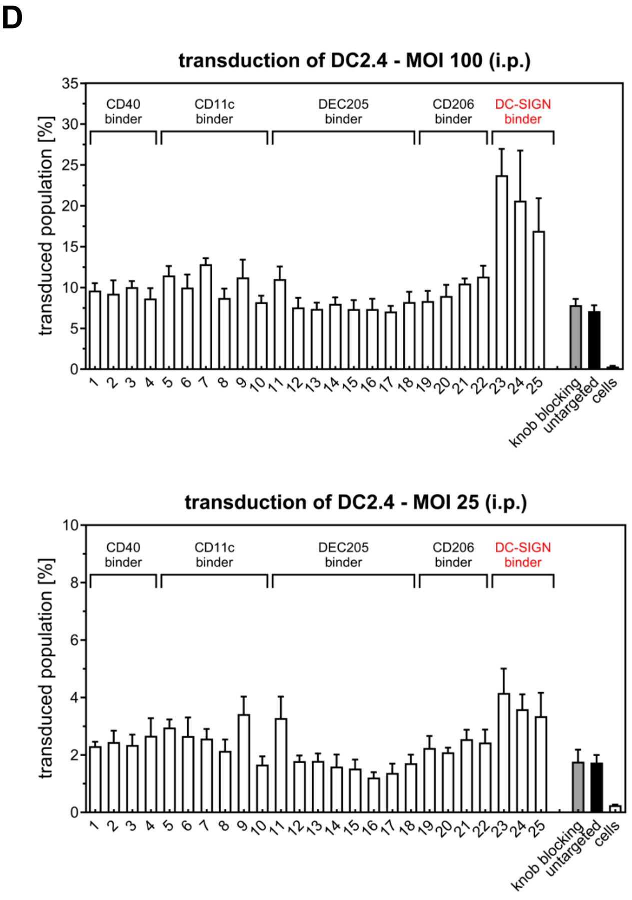

B

| nr | target | binder | format | reference | term |
|---|---|---|---|---|---|
| 1 | CD40 | sCD40L soluble murine | natural ligand | UniProt P27548 | |
| 2 | CD40 | R9 | scFv | Zhang et al. (2017) | |
| 3 | CD40 | R33 | scFv | Zhang et al. (2017) | |
| 4 | CD40 | 19-21 | scFv | Zhang et al. (2017) | |
| 5 | CD11c | N418 | scFv | Britton and Demangel (2004) | |
| 6 | CD11c | p17 cycl | peptide | Faham and Altin (2011) | |
| 7 | CD11c | p17 | peptide | Faham and Altin (2011) | |
| 8 | CD11c | p18 cycl | peptide | Faham and Altin (2011) | |
| 9 | CD11c | p18 | peptide | Faham and Altin (2011) | |
| 10 | CD11c | p30 | peptide | Faham and Altin (2011) | |
| 11 | DEC205 | NLDC | scFv | Britton and Demangel (2004) | |
| 12 | DEC205 | scFv12 | scFv | Dighe and Sinha (2020) | |
| 13 | DEC205 | scFv37 | scFv | Dighe and Sinha (2020) | |
| 14 | DEC205 | scFv39 | scFv | Dighe and Sinha (2020) | |
| 15 | DEC205 | scFv42 | scFv | Dighe and Sinha (2020) | |
| 16 | DEC205 | scFv83 | scFv | Dighe and Sinha (2020) | |
| 17 | DEC205 | scFv95 | scFv | Dighe and Sinha (2020) | |
| 18 | CD206 | RP-182 | peptide | Jaynes et al. (2020) | |
| 19 | CD206 | RP-185 | peptide | Jaynes et al. (2020) | |
| 20 | CD206 | RP-426 | peptide | Jaynes et al. (2020) | |
| 21 | CD206 | mUNO | peptide | Jaynes et al. (2020) | |
| 22 | CD206 | UNO | peptide | Jaynes et al. (2020) | |
| 23 | DC-SIGN | LASV GP1(M192R) | viral glycoprotein | UniProt P08669 | **$GP1_{M192R}$-adapter** |
| 24 | DC-SIGN | LASV GP1(M192R)/GP2 | viral glycoprotein | UniProt P08669 | **$GP1_{M192R}$-GP2-adapter** |
| 25 | DC-SIGN | LASV GP1 wt | viral glycoprotein | UniProt P08669 | **GP1 wt-adapter** |

C

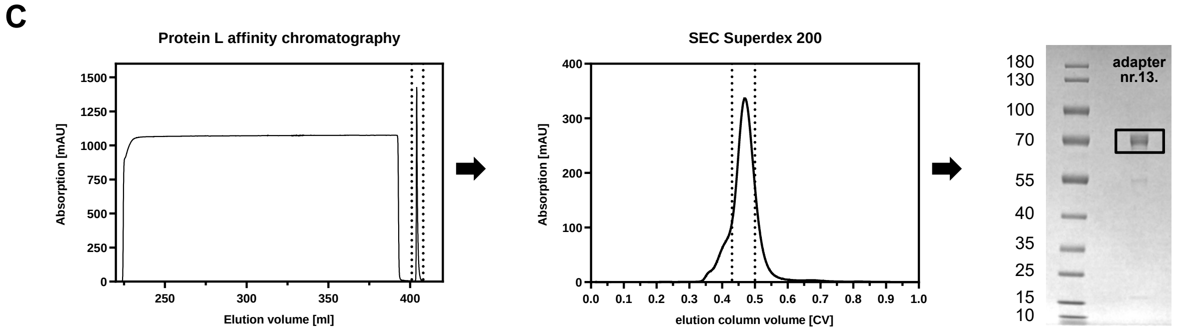

### Figure S2: Screening of retargeting adapters in mono-adapter format

(A) Schematic representation of a trimeric retargeting adapter. The retargeting module (grey) is fused by a flexible linker to the knob binding DARPin 1D3nc (green) and to the C-terminal trimerizing protein SHP, which leads to the formation of a stable clamp formed around the adenovirus fiber knob, also blocking the epitope for binding to CAR (coxsackievirus and adenovirus receptor). The retargeting domain redirects binding to a selected surface marker (target) and can mediate transduction to dendritic cells. (B) Table of tested retargeting adapters binding to CD40, CD11c, DEC205, CD206 and DC-SIGN. (C) Purification steps and quality control. Representative data of mono-adapter Nr. 13. The adapter was purified from CHO-S supernatants by Protein L affinity chromatography. Pooled elution fractions (dotted lines) were further purified by size exclusion chromatography (SEC) and the final pool (dotted lines) was stored in PBS at -20 °C. SDS-PAGE of the reduced sample confirmed high purity (calculated MW value is 57 kDa). (D) Murine cell line DC2.4 transduced by HAdV5$^{\Delta HVR7}$ encoding tdTomato. Transduced cell populations were quantified by flow cytometry (tdTomato+). Certain mono-adapters mediate transduction at the two MOIs shown here (100 or 25). As negative controls, HAdV5$^{\Delta HVR7}$ with blocked knob (E2_5 blocking adapter) and an untargeted control (natural transduction rate) were tested.

A

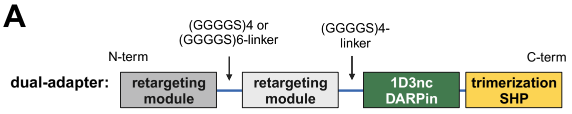

B

| nr | target | binder | linker | target | binder |
|---|---|---|---|---|---|
| 7 | - | - | - | CD11c | p17 |
| 10 | - | - | - | CD11c | p30 |
| 13 | - | - | - | DEC205 | scFv37 |
| 14 | - | - | - | DEC205 | scFv39 |
| 15 | - | - | - | DEC205 | scFv42 |
| 26 | DEC205 | scFv42 | (GGGGS)6 | CD11c | p17 |
| 27 | DEC205 | scFv42 | (GGGGS)4 | CD11c | p17 |
| 28 | DEC205 | scFv42 | (GGGGS)6 | CD11c | p30 |
| 29 | DEC205 | scFv42 | (GGGGS)4 | CD11c | p30 |
| 30 | CD11c | p17 | (GGGGS)6 | DEC205 | scFv42 |
| 31 | CD11c | p17 | (GGGGS)4 | DEC205 | scFv42 |
| 32 | CD11c | p30 | (GGGGS)6 | DEC205 | scFv42 |
| 33 | CD11c | p30 | (GGGGS)4 | DEC205 | scFv42 |
| 34 | CD11c | iCAM-1 D4loop | (GGGGS)6 | DEC205 | scFv42 |
| 35 | CD11c | iCAM-1 D4loop | (GGGGS)4 | DEC205 | scFv42 |
| 36 | CD11c | p17 | (GGGGS)4 | CD11c | p17 |
| 37 | CD11c | p30 | (GGGGS)4 | DEC205 | scFv37 |
| 38 | CD11c | p30 | (GGGGS)4 | DEC205 | scFv39 |
| 39 | CD11c | iCAM-1 D4loop | (GGGGS)4 | DEC205 | scFv37 |
| 40 | CD11c | iCAM-1 D4loop | (GGGGS)4 | DEC205 | scFv39 |

D

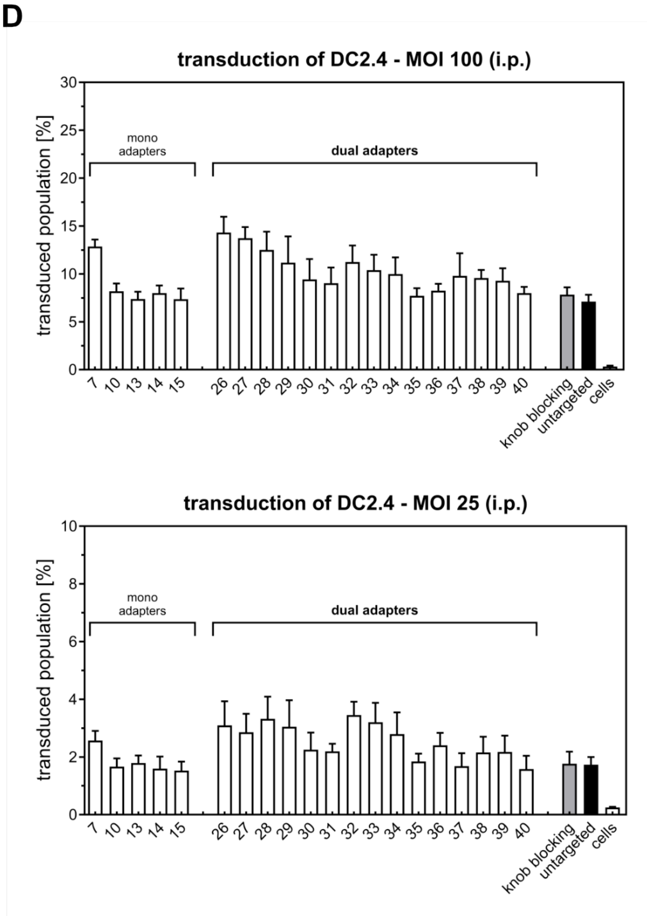

D

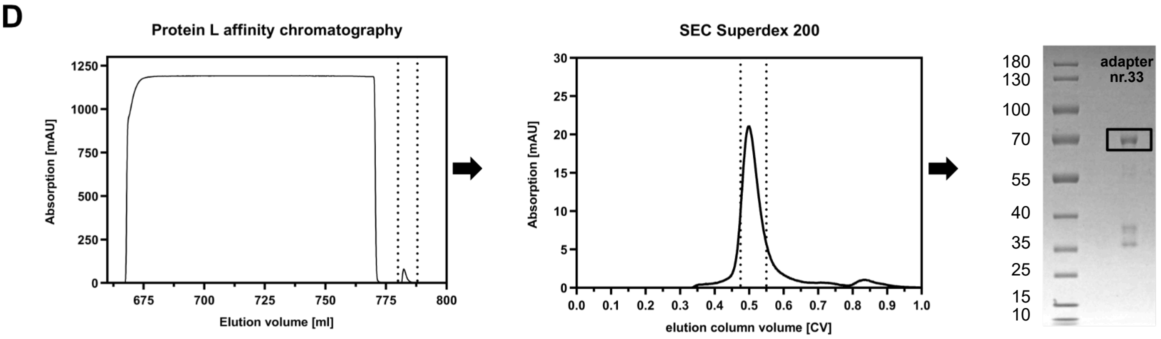

**Figure S3: Screening of retargeting adapters in dual-adapter format targeting DEC-205 and CDllc**

(A) Schematic representation of a trimeric dual retargeting adapter. The two retargeting modules (grey), separated by a $(GGGGS)_4$ or $(GGGGS)_6$-linker, are fused by a flexible linker to the knob-binding DARPin 1D3nc (green) and to the C-terminal trimerizing protein SHP, which leads to the formation of a stable clamp formed around the adenovirus fiber knob, also blocking the epitope for binding to CAR (coxsackievirus and adenovirus receptor). The two retargeting domains redirect binding to selected surface markers (targets) and can mediate transduction to dendritic cells. (B) Table of tested dual-retargeting adapters binding to DEC-205 and CD11c with the corresponding mono-adapters as control. (C) Purification steps and quality control. Representative data of dual-adapter Nr. 33. The adapter was purified from CHO-S supernatants by Protein L affinity chromatography. Pooled elution fractions (dotted lines) were further purified by size exclusion chromatography (SEC) and the final pool (dotted lines) was stored in PBS at -20 °C. SDS-PAGE of the reduced sample confirmed purity (calculated MW value is 61 kDa) (D) Murine cell line DC2.4 transduced by $HAdV^{\Delta HVR7}$ encoding tdTomato. Transduced cell populations were quantified by flow cytometry (tdTomato+). Retargeting modules in mono-adapter format mediate only low transduction levels. In contrast, certain module combinations in the dual-adapter format targeting CD11c and DEC205 show synergistic effects and rescue adapter-mediated transduction. As negative controls, $HAdV^{\Delta HVR7}$ with blocked knob (E2_5 blocking adapter) and an untargeted control (natural transduction rate) were tested.

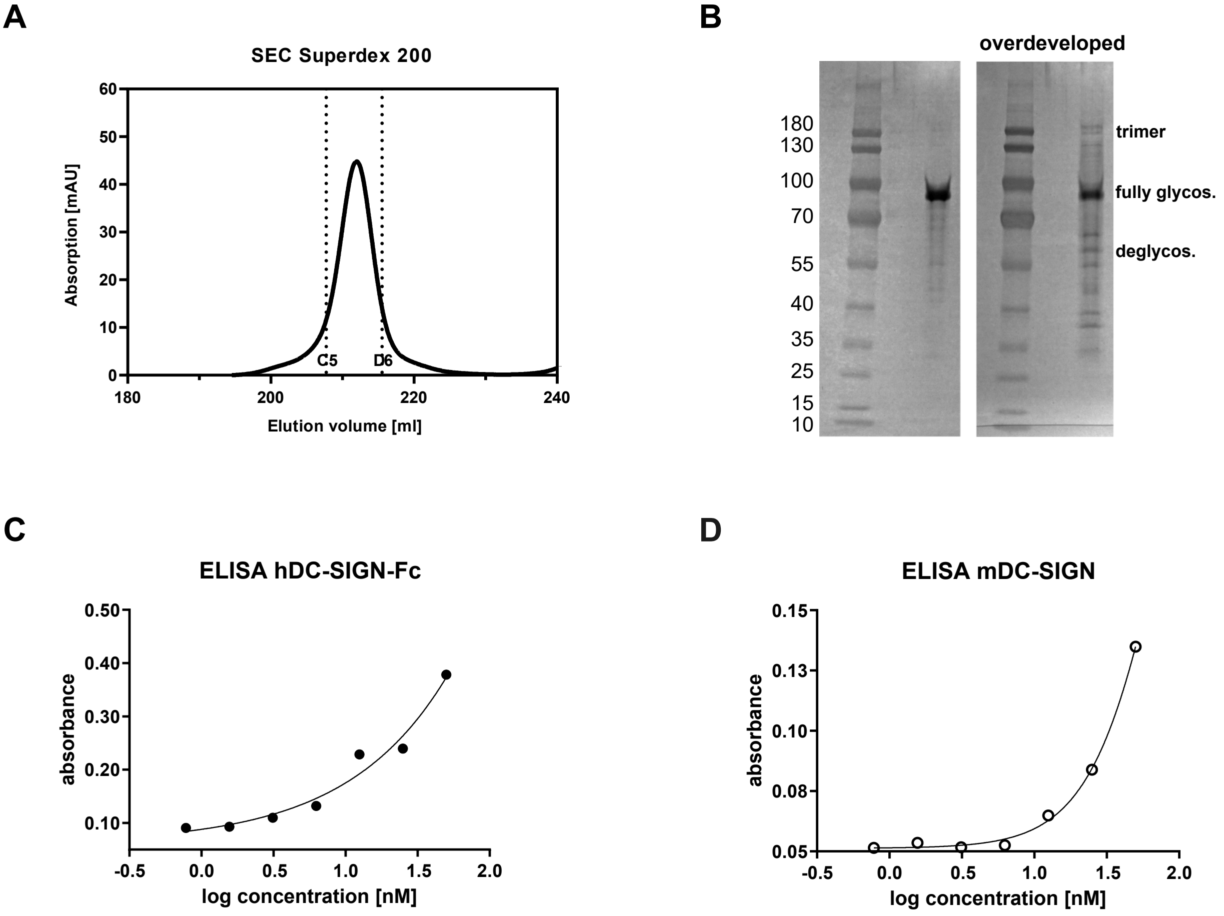

**Figure S4: Quality control and characterization of α-DC adapter**

(A) α-DC adapter protein, purified by ion exchange chromatography, was polished by size exclusion chromatography (SEC, Superdex 200). Elution profile of size exclusion chromatography with depicted pooled fractions (C5-D6) demonstrates that the adapter trimer is monodisperse. (B) SDS-PAGE and detection by silver stain of reduced adapter protein. Adapter monomer with a calculated molecular weight of 52 kDa runs at an apparent molecular weight of ~80 kDa due to seven N-linked glycans of GP1(LASV). SDS-PAGE (left) shows high protein purity. SDS-PAGE (right) of the same sample, this time with overdeveloped silver staining, reveals traces of adapter trimer (~240 kDa), and a pattern of deglycosylated adapter (55-70 kDa) next to the fully glycosylated adapter (~80 kDa) representing the main fraction. (C) Binding of coated adapter to human DC-SIGN-Fc by ELISA. After coating of adapter protein (20 nM), serial dilutions of human DC-SIGN-Fc (SinoBiological, 10200-H01H) were added. Binding was detected via Fc (anti human Fc-HRP labelled antibody). Antibodies are listed in the Materials and Methods section. (D) Binding of coated adapter to mouse DC-SIGN by ELISA. After coating of adapter protein (20 nM), serial dilutions of mouse DC-SIGN (R&D Systems, 8345-DC) were added. Bound DC-SIGN was detected with primary anti-mouse DC-SIGN mouse antibody and secondary anti-mouse IgG-HRP-labeled antibody. Antibodies are listed in the Materials and Methods section.

## A

**iCAM1 D4 loop**

Identity: 66.7%
Similarity: 66.7%

```
human  P05362  404  P G N W T W P E N S Q Q T P M  418
mouse  P13597  407  L G N W T W Q E G S Q Q T L K  421
```

## B

**CD11c**

Identity: 67.6%
Similarity: 83.0%

```
human  P20702  149  E Q D I V F L I D G S G S I S S R N F A T M M N F V R A V I S Q F Q R P S T Q F S L M Q F S N K F Q  198
mouse  Q9QXH4  150  D Q D I V F L I D G S G S I S S T D F E K M L D F V K A V M S Q L Q R P S T R F S L M Q F S D Y F R  199

                199  T H F T F E E F R R S S N P L S L L A S V H Q L Q G F T Y T A T A I Q N V V H R L F H A S Y G A R R  248
                200  V H F T F N N F I S T S S P L S L L G S V R Q L R G Y T Y T A S A I K H V I T E L F T T Q S G A R Q  249

                249  D A A K I L I V I T D G K K E G D S L D Y K D V I P M A D A A G I I R Y A I G V G L A F Q N R N S W  298
                250  D A T K V L I V I T D G R K Q G D N L S Y D S V I P M A E A A S I I R Y A I G V G K A F Y N E H S K  299

                299  K E L N D I A S K P S Q E H I F K V E D F D A L K D I Q N Q L K E K I F A I  336
                300  Q E L K A I A S M P S H E Y V F S V E N F D A L K D I E N Q L K E K I F A I  337
```

## C

**DC-SIGN**

Identity: 64.0%
Similarity: 77.2%

```
human  Q9NNX6  253  C H P C P W E W T F F Q G N C Y F M S N S Q R N W H D S I T A C K E V G A Q L V V I K S A E E Q N F  302
mouse  Q91ZX1  105  C R S C P W D W T H F Q G S C Y F F S V A Q K S W N D S A T A C H N V G A Q L V V I K S D W W Q N F  154

                303  L Q L Q S S R S N R F T W M G L S D L N Q E G T W Q W V D G S P L L P S F K Q Y W N R G E P N N V G  352
                155  L Q   Q T S K K R G Y T W M G L I D M S K E S T W Y W V D G S P L T L S F M K Y W S K G E P N N L G  203

                353  E E D C A E F S G N G W N D D K C N L A K F W I C K K S A A S C R D E  388
                204  E E D C A E F R D D G W N D T K C T N K K F W I C K K L S T S C P S K  238
```

**Figure S5: Cross-species binding of α-DC adapter is supported by human and mouse sequence similarities**

Pairwise sequence alignment (EMBOSS Needle) of relevant UniProt entries. (A) The peptide of the iCAM1 D4 loop (used as CD11c targeting module of the α-DC adapter) reveals high sequence similarity between the human and murine protein sequence. This high sequence similarity explains binding of the iCAM1 D4 loop to CD11c on both human and murine cells. The α-DC adapter was chosen to contain the murine iCAM1 D4 loop to achieve high efficiency in the *in vivo* mouse model. (B) High sequence similarity between human and murine ligand binding ecto domain αI of CD11c. High sequence similarity supports cross-specific binding of murine iCAM1 D4 loop to CD11c (C) High sequence similarity between human and murine C-terminal carbohydrate-binding domain of DC-SIGN. High sequence similarity supports cross-specific binding of glycosylated GP1(LASV) module targeting DC-SIGN.

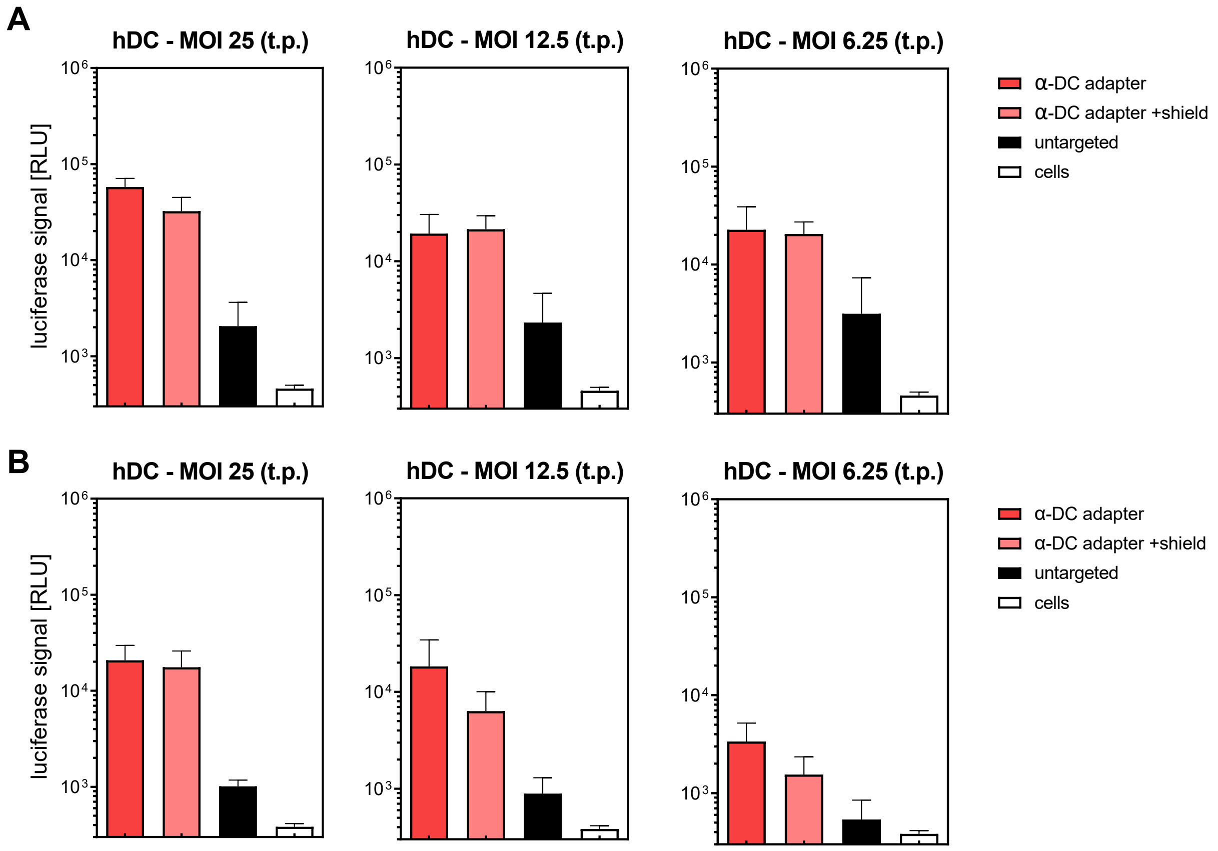

**Figure S6: Adapter mediates HAdV5 transduction in human mo-DC**

Complementary transduction data with additional human donors. (A) CD14$^+$ monocytes of donor B were differentiated to mo-DCs. Generated DCs of donor B were then transduced with HAdV5 encoding ffLuciferase. Two days post infection, luciferase activity of the cell lysate was measured. The targeted adapter enhances transduction efficiency. (B) The same procedure was carried out starting with CD14$^+$ monocytes of donor C. t.p.: transducing particles

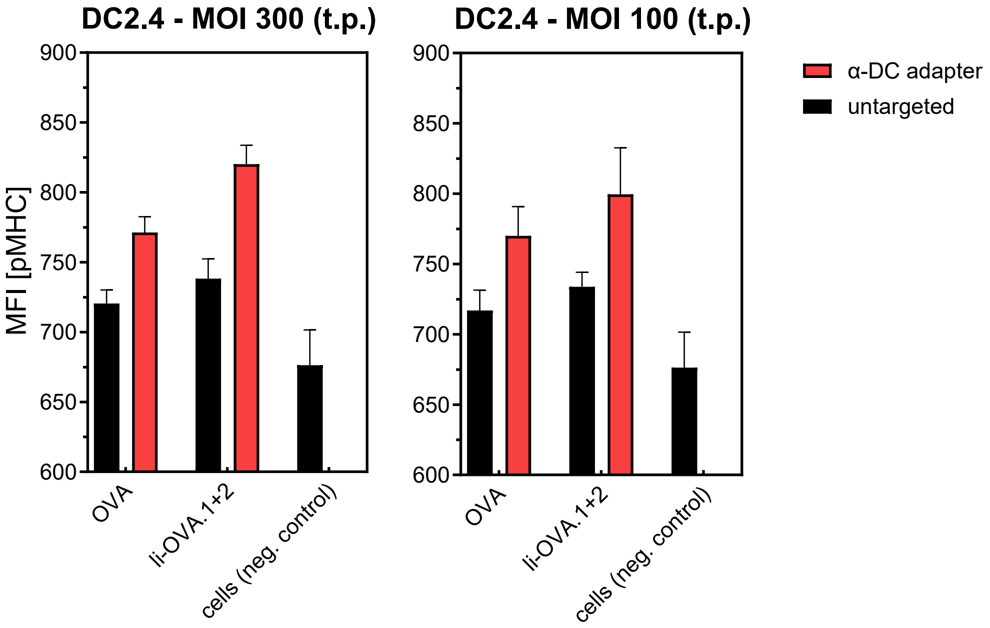

**Figure S7: Invariant chain modulates MHC class I antigen presentation**

Transduction of DC2.4 cells with HAdV5 (MOI 300 or 100) encoding wild-type ovalbumin or truncated ovalbumin fusion to invariant chain. Two days post infection pMHCI was detected (via SIINFEKL antigen presentation) by flow cytometry. The median fluorescence intensity of the total single-cell population is shown. Antigen trafficking to the MHC class I (and MHC class Il) loading compartment by invariant chain fusion improved MHC class I antigen presentation in comparison to the wild-type ovalbumin transgene. Populations of targeted HAdV5 showed increased presentation in comparison to untargeted control, due to the increased number of transduced cells. t.p.: transducing particles

A

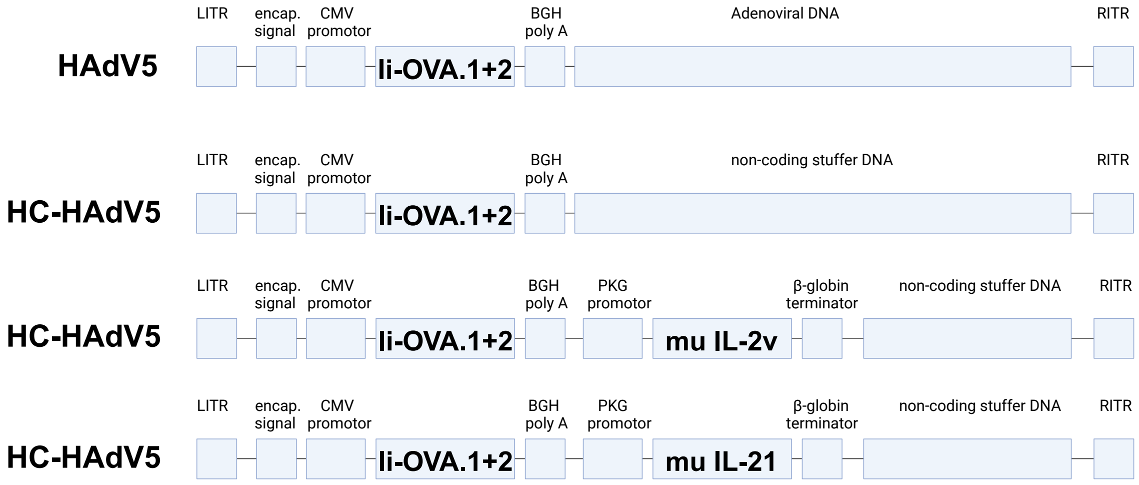

B

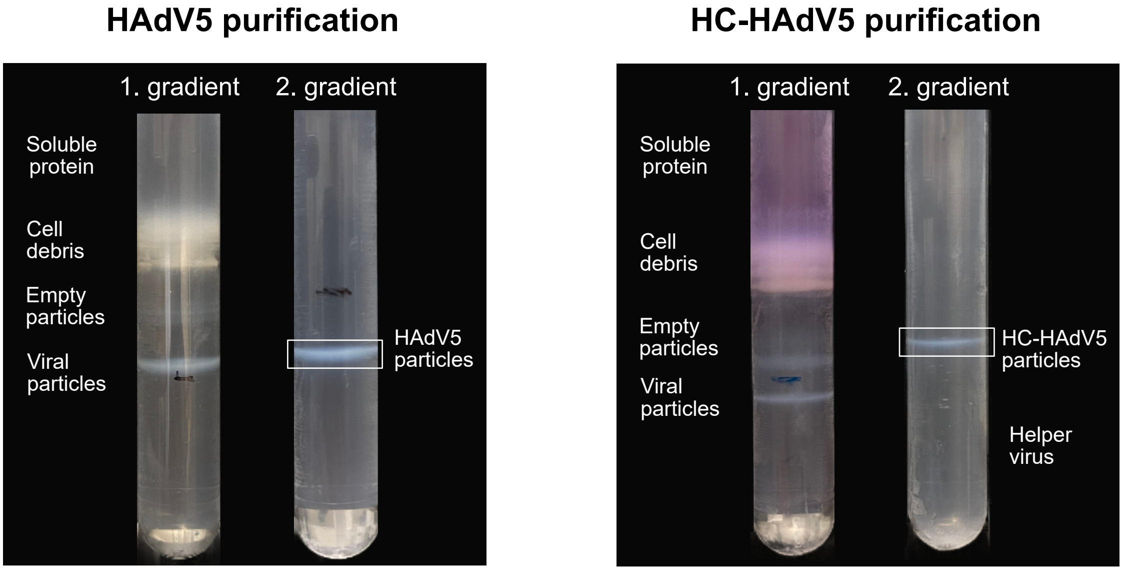

C

| Transgene | HAdV5/µL |
|---|---|
| Ii-OVA1+2 | $1.44 \times 10^8$ |

| Transgene | HC-HAdV5/µL | HV particles/µL | HV cont. [%] |
|---|---|---|---|
| Ii-OVA1+2 | $1.65 \times 10^7$ | $3.44 \times 10^4$ | 0.21 |
| Ii-OVA1+2 +IL-2v | $1.97 \times 10^7$ | $6.64 \times 10^4$ | 0.34 |
| Ii-OVA1+2 +IL-21 | $1.85 \times 10^7$ | $1.54 \times 10^5$ | 0.83 |

**Figure S8: Vector production and purification**

(A) Representative schematic vector map of produced viral particles (not to scale). The antigen Ii-OVA1+2 is encoded under the CMV promotor. The co-stimulatory cytokines (IL-2v or IL-21) are encoded under the PGK promotor. The first-generation HAdV5 vector (ΔE1, ΔE3) encodes adenoviral genes and is amplified in HEK293 cells (which express E1). HC-AdV5 with all adenoviral genes deleted is amplified in cell line 116 with a helper virus (not shown), which expresses all AdV genes except E1, but with a removable encapsulation signal. All vectors are flanked by inverted terminal repeats, important for DNA replication. The encapsulation signal is important for packaging viral DNA into the formed viral particle. (B) Two-step CsCl gradient for purification of viral particles from producer cell lysate supernatant. In a first gradient, viral particles are separated from empty particles, cell debris and soluble proteins. From the isolated band, viral particles are further purified in a second gradient. For HC-HAdV5, helper virus is separated during the second gradient. (C) HAdV5 and HC-HAdV5 are produced in high quantity and purity. For HC-HAdV5, only very low helper virus impurities were detected. All numbers depicted represent transducing particles quantified by qPCR.

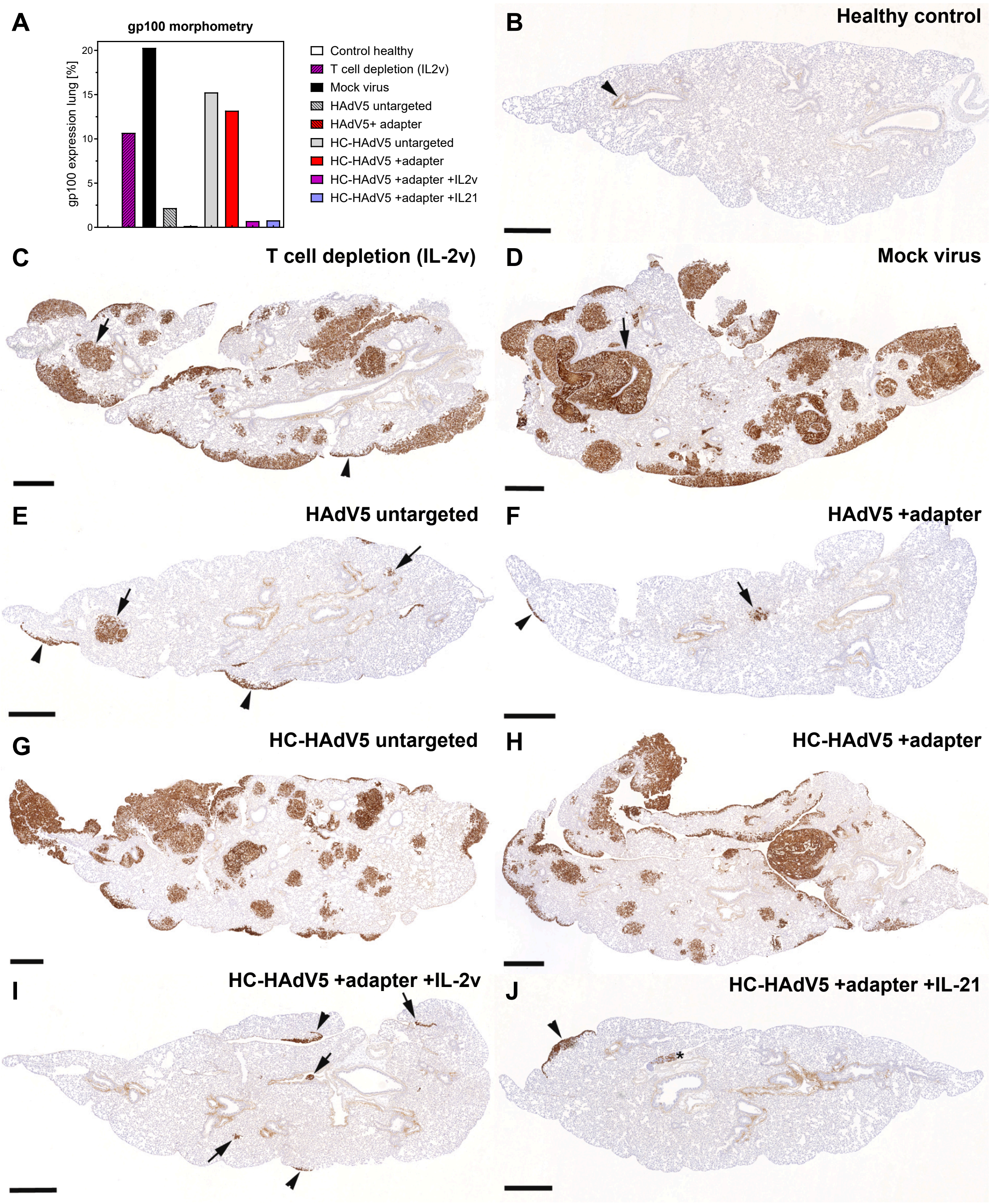

**Figure S9: Immunohistochemical staining of lung sections for the melanoma marker gp100**

Representative lung sections from one animal of each treatment group. (A) Quantitative staining analysis of gp100, n=1. Percentage calculated by immuno-stained area divided by total area × 100. (B) The lung of the control animal is free of any metastasis. Arrowhead: non-specific staining of bronchiolar smooth muscle layer, as seen in all stained lungs (discarded in the morphometric analysis). (C) The lung of the T cell-depleted (expressing li-OVA.1+2 and IL-2v) mouse exhibits numerous metastatic foci, in both subpleural (arrowhead) and parenchymal (arrow) locations. (D) The lung of the mock-treated mouse exhibits numerous subpleural and parenchymal metastatic foci, often pronounced around vessels (arrow). (E) In the mouse treated with untargeted HAdV5 (expressing li-OVA.1+2), a few small subpleural (arrowheads) and parenchymal (arrows) metastatic foci are observed. (F) The lung sections of the mouse treated with HAdV5 +adapter (expressing li-OVA.1+2) exhibits each one subpleural (arrowhead) and parenchymal (arrow) metastatic focus. (G) After treatment with untargeted HC-HAdV5 (expressing li-OVA.1+2), numerous variably sized metastatic foci are seen. (H) Also the lung of a mouse treated with HC-HAdV5 +adapter (expressing li-OVA.1+2) harbors numerous variably sized metastatic foci. (I) After treatment with HC-HAdV5 +adapter +IL-2v there are only a few very small subpleural (arrowheads) and parenchymal (arrows) metastatic patches. (J) Treatment with HC-HAdV5 +adapter +IL-21 also only results in rare metastatic foci (arrow: subpleural metastasis). The bronchial lymph node (*) harbors gp100-positive neoplastic cells, a feature that was also seen in other mice. Immunohistochemistry for gp100, hematoxylin counterstain. Bars = 500 µm.

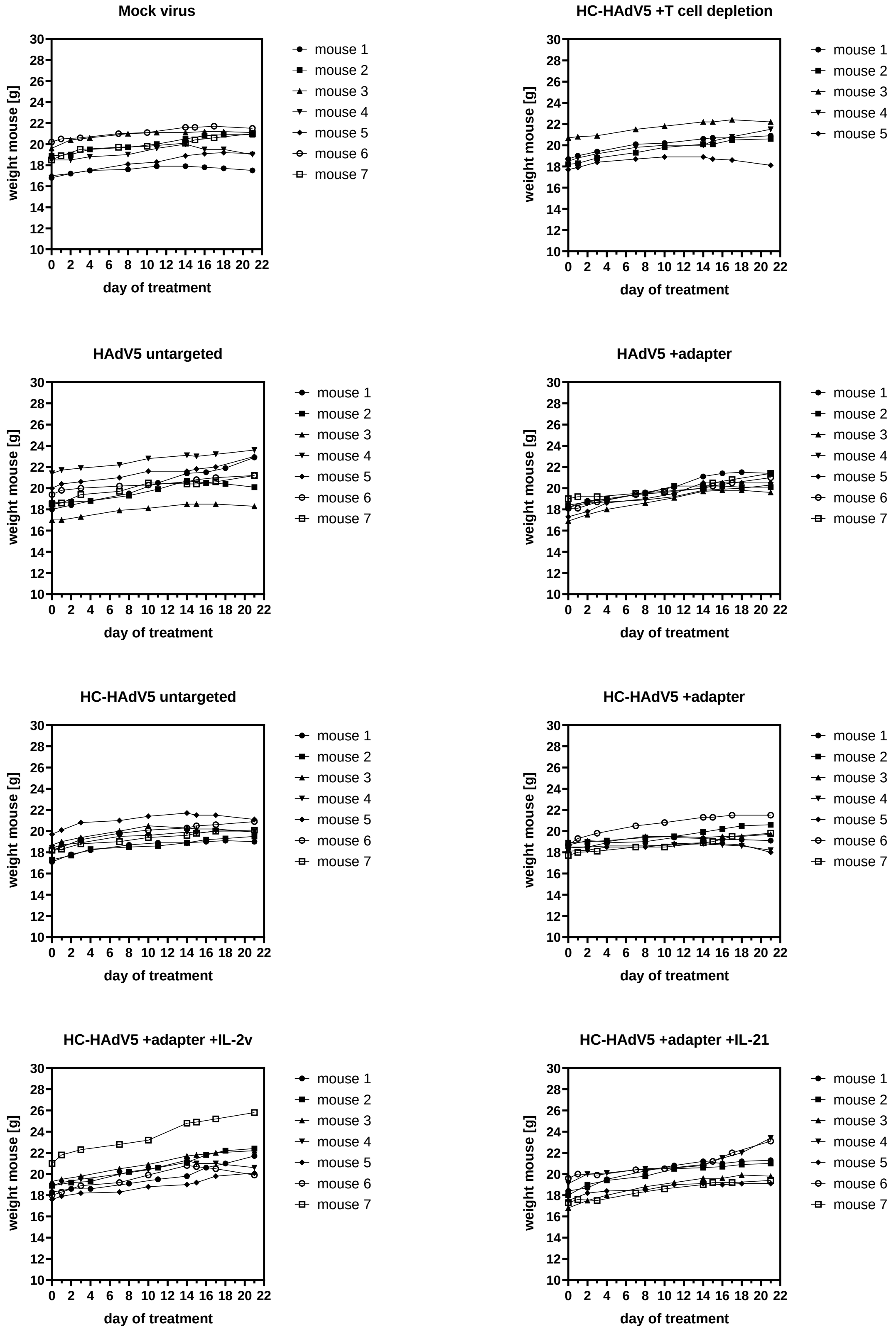

**Figure S10: Acute toxicity monitored by body weight. Tumor vaccination study $1\times10^7$ vp/injection**

The body weight of each mouse in every treatment group was measured during the course of the experiment with the lower virus dose. No significant loss of body weight (> 5%) was observed. Every mouse had stable or increasing body weight during the treatment.

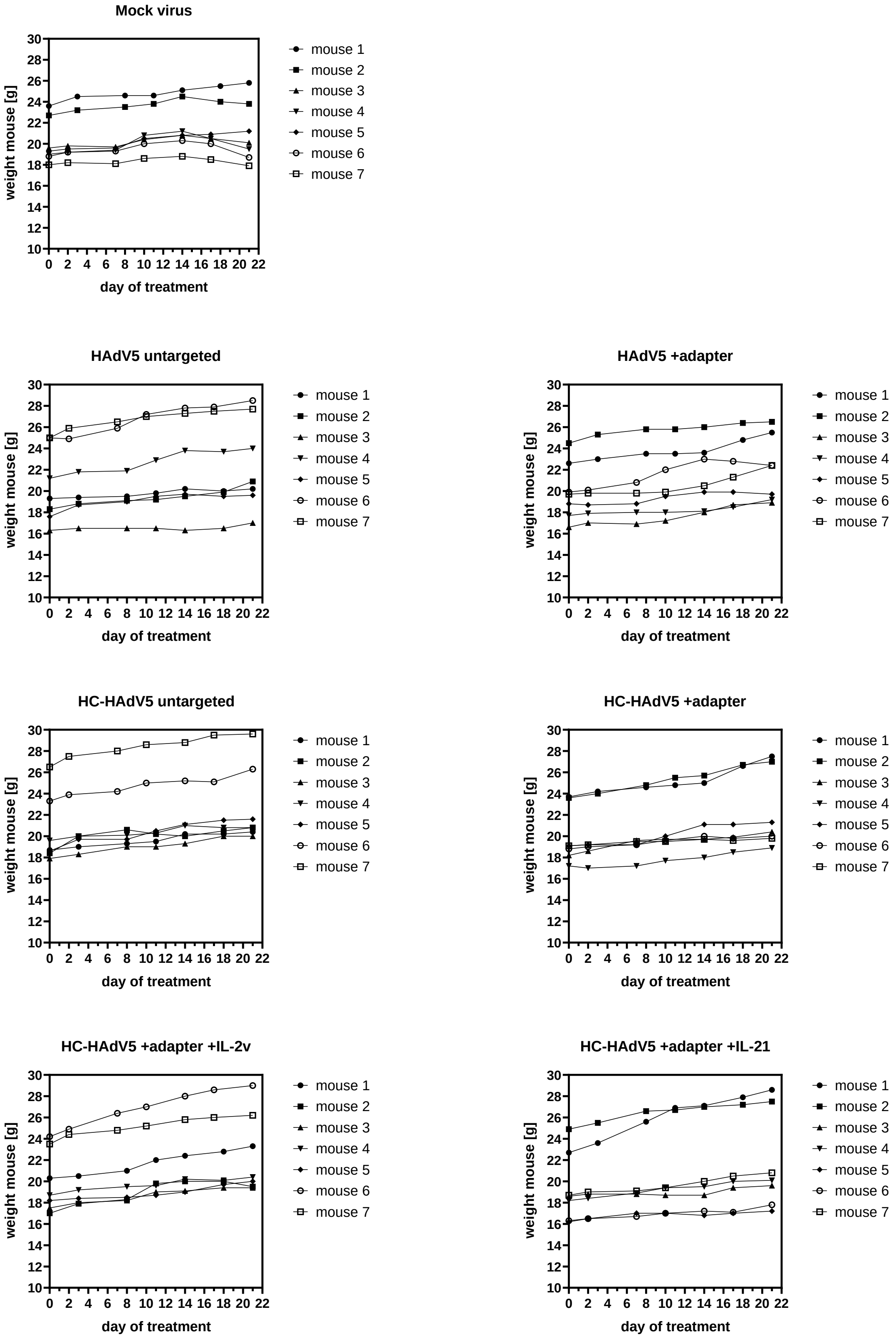

**Figure S11: Acute toxicity monitored by body weight. Tumor vaccination study $3\times10^8$ vp/injection**

The body weight of each mouse in every treatment group was measured during the course of the experiment with the higher virus dose. No significant loss of body weight (> 5%) was observed. Every mouse had stable or increasing body weight during the treatment.
